# Transcriptomic analysis of repeat expansion-ataxias uncovers distinct non-neuronal cell type-specific signatures of disease across the human brain

**DOI:** 10.1101/2025.01.03.631007

**Authors:** Zhongbo Chen, Amy Hicks, Jonathan Brenton, Melissa Grant-Peters, Regina H. Reynolds, Emil K. Gustavsson, Clarissa Rocca, Guillermo Rocamora-Perez, Raquel Garza, Sonia Garcia-Ruiz, Natalia Dominik, Claire Anderson, Toby Curless, Kylie Montgomery, Hannah Macpherson, Suran Nethisinghe, Daria Gavriouchkina, Modesta Blunskyte-Hendley, Aine Fairbrother-Browne, Jasmaine Lee, Huihui Luo, Stephanie Efthymiou, David Murphy, Fairlie Hinton, Juan Botia, Andrea Cortese, Nicholas Wood, Paola Giunti, John Hardy, Johan Jakobsson, Sonia Gandhi, Arianna Tucci, Catriona McLean, Zane Jaunmuktane, Henry Houlden, Mina Ryten

## Abstract

Hereditary ataxias are a heterogeneous group of neurogenetic conditions characterised by the clinical syndrome of progressive loss of coordination from neurodegeneration of the cerebellum. A commonality across the most prevalent ataxias is the underlying disease mechanism secondary to expansions of short tandem DNA repeats. There is currently an incomplete understanding of the pathogenic mechanisms of these repeat expansion disorders, a core feature of which revolves around RNA-dysregulation. In this study, we used both bulk and single nuclear RNA-sequencing to study post-mortem brain tissue of human donors with a range of repeat-expansion ataxias to reveal further mechanistic insights.

We compared post-mortem paired cerebellar and frontal cortex tissue bulk RNA-sequencing data from 23 ataxia patients and 22 sex-, age-matched controls from two brain banks (spinocerebellar ataxia (SCA)1, SCA2, SCA6, SCA7, SCA17, Friedreich’s ataxia (FRDA), and 7 cases with unknown molecular diagnoses). We analysed bulk RNA-sequencing data for transcript usage, differential and cell-type-specific expression to transcriptomically profile these diseases. We also generated single nuclear RNA-sequencing data of the cerebellum from donors with SCA1, SCA2, SCA6 and FRDA to decipher changes in cell type proportions in the disease state.

Using this approach, we found that: (i) despite the commonalities in the genetics of ataxia, there were components of their transcriptional signatures which were distinct; (ii) there were extensive transcriptional changes evident not only in the cerebellum but also the frontal cortex in ataxia cases; (iii) activation of immune and inflammatory pathways, as well as involvement of non-neuronal cell types was a feature of all ataxias to a lesser or greater extent.

This study provides a novel resource to understand the mechanisms of disease in ataxia. Furthermore, taken together, these results highlight immune pathways and the role of non-neuronal cell types as early and potentially important therapeutic targets. These findings provide a map of transcriptomic changes in ataxia to further understanding of the underlying pathogenesis.

## Introduction

Hereditary ataxias are a group of neurodegenerative disorders centred on the progressive degeneration of the cerebellum and its connections^1–3^. While they are clinically and genetically heterogeneous, collectively they affect 10 in every 100,000 people^4–6^. Individuals with ataxia suffer from the predominant clinical syndrome of debilitating incoordination and loss of balance. Despite ataxias being forerunners for precision medicine given their defined genetic aetiology, limited therapeutic options currently exist. This is due in large part to the incomplete understanding of their underlying disease biology.

Beyond their clinical commonality, the most prevalent subtypes of ataxia share the single underlying pathogenic mechanism of repeat expansion^7,8^. Here, short tandem repeat (STR) sequences up to six bases in tandem, are repeated multiple times to cause disease^7,8^. Repeat expansions account for a high proportion of spinocerebellar ataxia (SCA) disorders explaining three quarters of all molecularly diagnosed autosomal dominant cases globally^1^, and the most common autosomal recessive ataxia in Europeans, namely Friedreich’s ataxia (FRDA)^9^. Strikingly, more than half of the 60 or so established monogenic repeat expansion disorders are associated with ataxia, yet we still do not know why the cerebellum is particularly vulnerable^7^. Recent progress in the field leading to diagnosis of previously unsolved late-onset ataxia cases has generated much interest^10–16^. For example, the discovery of novel repeat expansions such as the exonic *ZFHX3-*GGC expansion underlying late-onset SCA4 has resolved a 25-year diagnostic conundrum, and defined a novel disease entity of polyglycine disorders^10,11,15,16^.

Conventionally, while it was proposed that disease in exonic CAG-repeat-associated SCAs is mediated by toxic gain-of-function from canonically translated polyglutamine protein, it is now appreciated that multiple disease mechanisms can coexist even within one repeat disorder let alone across disorders^1,17^. For example, in SCA2, RNA-binding protein (RBP) sequestration from the repeat-transcribed RNA and repeat-associated non-AUG (RAN) translation have been shown to occur in addition to toxic polyglutamine accumulation^18^. It is now appreciable that there are gaps in understanding the putative disease mechanisms (**Supplementary Table 1** summarises unknown mechanisms of disease along with clinical features of the studied diseases). Remarkably, most of the causative genes in the SCAs involve transcriptional regulation or RNA-processing^19^ and RNA dysregulation is a key driver of pathogenesis^20^. Therefore, transcriptome-wide changes would be expected to be central to the pathogenesis, making the delineation of the disease at the RNA level crucial.

While previous studies have focused on either mouse or patient-derived cell line data, pathogenesis in human brain tissue remains largely unresolved. Recent key advances in Huntington’s disease using human postmortem brain tissue suggested that somatic repeat expansion is a key pathogenic driver of transcriptome-wide changes in a cell-type specific manner^21,22^, a finding that has been difficult to recapitulate in animal or cell line models. This highlights the importance of studying disease using human data but a major barrier has been the availability of adequate tissues available in these rare disorders. A handful of transcriptomic studies in ataxia using human post-mortem brain samples are emerging, revealing dysregulation and involvement of non-neuronal cell types in fragile X-associated tremor-ataxia syndrome (FXTAS)^23^ and SCA1^24^. A separate study using ataxia telangiectasia human cerebellar tissue proposed a role for LINE1 activation^25^. Furthermore, Purkinje cell enriched RNA-sequencing in SCA7 have shed light on specific transcriptional changes in the disease state^26^. Despite this paucity of data in transcriptomic analysis of hereditary ataxia, RNA-based therapies are currently in development for SCAs^27^. Some promise has been seen in *ATXN2* mRNA lowering anti-sense oligonucleotide (ASO) treatment in SCA2 mouse models^28^. Thus, the promise brought by RNA-based therapeutics in neurological diseases make it ever more pressing to capture an accurate transcriptomic map of the ataxia disease state.

The aim of this study was to: (i) determine whether all SCAs have a common core transcriptomic signature which only varies relative to their symptom severity, and (ii) determine if there is evidence for a loss of RNA regulatory functions. Leveraging bulk-tissue derived from two brain regions (cerebellum and frontal cortex) and snRNA-sequencing data from the cerebellum of individuals across six different molecularly diagnosed ataxias (SCA 1, 2, 6, 7, 17 and FRDA) and those without molecularly confirmed forms of ataxia despite standardised diagnostic approaches, we characterised the landscape of disease through transcriptome-wide differential gene expression and splicing. Through this process, we uncovered distinct disease pathways, unexpected immune activation and contribution from glial cells in the disease process across all ataxias, improving the mechanistic understanding of pathogenesis.

## Methods

### Sample selection

Given the heterogeneity of hereditary ataxias and the paucity of post-mortem cases available compared with other neurodegenerative disorders, cases were selected to reflect the most common ataxia presentations. W e included seven as-yet molecularly undiagnosed cases of hereditary ataxia, as defined by cases in which standardised clinical testing as used in the NHS Genomic Medicine Service did not yield any pathogenic variants, to identify any transcriptomic signatures underlying a common clinical presentation. Control individuals were defined by a lack of history of neurological disease or presentations, and no definitive pathological diagnoses. Control individuals were age-, sex-, and brain bank-matched to cases. Post-mortem brain tissue was obtained from two brain banks only, the Queen Square Brain Bank (QSBB, UK) and the Victorian Brain Bank (VBB, Australia), to minimise inter-brain bank sample variability. For each individual, both cerebellar and frontal cortex tissue were used to study pathology across the brain regions, with the hypothesis being that the cerebellar cortex would be the region most affected in ataxia.

A total of 91 samples were analysed (46 from cases and 45 from controls: **Table 1**, full anonymised details available in **Supplementary Table 2**). In total, we used paired cerebellar and frontal cortex tissue from: 4 individuals with FRDA; 3 individuals with SCA1; 3 individuals with SCA2; 3 individuals with SCA6; 2 individuals with SCA17; 1 individual with SCA7 and 7 individuals who had molecularly undiagnosed ataxia.

**Table 1.** Total number of samples analysed. Number of samples (n) for each category are shown. QSBB refers to Queen Square Brain Bank and VBB refers to the Victorian Brain Bank. Sex (n male) refers to the number of samples derived from male individuals. FRDA refers to Friedreich’s ataxia, SCA refers to spinocerebellar ataxia.

| Sample | Total samples<br>(n) | Cerebellar samples<br>(n) | Frontal samples<br>(n) | QSBB samples<br>(n) | VBB samples<br>(n) | Sex (n male) |
| --- | --- | --- | --- | --- | --- | --- |
| Cases | 46 | 23 | 23 | 28 | 18 | 24 |
| Controls | 45 | 23 | 22 | 26 | 19 | 25 |
| **Cases by molecular diagnosis** |  |  |  |  |  |  |
| FRDA | 8 | 4 | 4 | 4 | 4 | 4 |
| SCA1 | 6 | 3 | 3 | 4 | 2 | 4 |
| SCA2 | 6 | 3 | 3 | 0 | 6 | 2 |
| SCA6 | 6 | 3 | 3 | 2 | 4 | 4 |
| SCA7 | 2 | 1 | 1 | 2 | 0 | 0 |
| SCA17 | 4 | 2 | 2 | 2 | 2 | 2 |
| Molecularly diagnosis unknown | 14 | 7 | 7 | 14 | 0 | 8 |

Statistical comparisons between the demographic variables were performed in R (v.4.0.5), using chi-squared test for categorical variables and Wilcoxon rank-sum test to compare continuous variables in a pairwise manner. FDR-correction for multiple testing was used where applicable. Correlations between covariates used Pearson’s correlation coefficient and for categorical variables, Kruskal Wallis test.

#### Ethics

A materials transfer agreement was signed between the host institution (UCL) and the two transferring brain banks for the supply of fresh frozen human brain samples for research purposes. Brain donors had been previously consented for donation by the recipient brain bank in accordance with local ethics committee approval. All tissues were handled and stored in line with the Human Tissue Act 2004. Ethical approval for genetics research in human brain tissue was under the remit of Project 07/N018 (UCLH).

#### Tissue preparation

All brain samples had been freshly frozen at post-mortem retrieval and stored in either QSBB or VBB. Two cerebellar cortex and two frontal cortex samples for each individual were chipped by a neuropathologist (ZJ) from QSBB to ensure consistency of sampling. The anterior superior frontal cortex and superior medial cerebellar cortex at the mid level of the dentate nucleus were sampled for all individuals to minimise anatomical variation. Each sample contained approximately 50 to 100 g of frozen tissue. Both cerebellar and frontal cortex tissue samples were sent for RNA extraction. One frontal cortex sample for each individual was used for DNA extraction for molecular confirmation.

### Confirmation of molecular diagnosis

DNA was extracted from the frontal cortex tissue for each individual using MagAttract HMW DNA Kit (Qiagen) following manufacturer’s instructions with resulting mean (SD) concentrations of 159.25 (111.15) ng/µL. The most common repeat expansions associated with hereditary ataxia were screened in sequential order (**Supplementary Figure 1**). First, samples were screened for the most common SCAs, as per Great Ormond Street Hospital NHS Diagnostic Lab testing protocol using tethered PCR. Positive samples with known and validated repeat expansions were used for each of the disorders screened as experimental controls. Allele sizes of the CAG repeats in *ATXN1* (SCA1), *ATXN2* (SCA2), *ATXN3* (SCA3), *CACNA1A* (SCA6) and *ATXN7* (SCA7) and CAG/CAA repeats of *TBP* (SCA17) were estimated using PCR. Samples were subsequently screened for the biallelic repeat expansion in *FXN* (FRDA) using flanking PCR and RP-PCR (**Supplementary Methods**). Samples negative for the above pathogenic expansions were then screened for more recently discovered repeat expansion disorders with protocols published as follows: *NOTCH2NLC* associated with neuronal intranuclear inclusion disease^29–31^); *RFC1* associated with CANVAS^14^ and *FGF14* associated with SCA27B^12^. Together, these ten repeat expansions account for a significant proportion of late adult-onset ataxia cases. Cases that tested negative for these ten repeat expansion disorders and had no pathogenic or likely pathogenic variants identified from whole-exome sequencing were categorised as molecularly undiagnosed.

#### RNA isolation

RNA extraction from brain tissue was carried out by BioXpedia A/S (Denmark). Samples were lysed with QIAzol and RNA extracted using the RNeasy 96 Kit (Qiagen) with an on-membrane DNase treatment, as per manufacturer instructions. Samples were quantified by absorption on the QIAxpert (Qiagen), and RNA integrity number (RIN) measured using the Agilent 4200 Tapestation (Agilent). The median RIN was 6.0 (interquartile range: 4.75 – 6.8) (**Supplementary Table 2**).

### Bulk-tissue RNA-sequencing data generation

UCL Genomics Facility constructed the bulk-tissue RNA-sequencing libraries and conducted the subsequent RNA-sequencing. For each sample, 500 ng of total RNA was used as input for cDNA library construction with the KAPA mRNA HyperPrep Kit (Roche), as per manufacturer instruction, with the aim of capturing poly-A-selected mRNA. mRNA was captured using magnetic oligo-dT beads and fragmented using heat and magnesium. The first strand cDNA synthesis used random priming, followed by combined second strand synthesis and A-tailing. Subsequently, adapter ligation was followed by library amplification in a strand-specific manner (further details in: https://elabdoc-prod.roche.com). To minimise read mis-assignment in downstream sample de-multiplexing, xGen Dual Index UMI Adapters (Integrated DNA Technologies, Inc.) were used. Libraries were multiplexed on the NovaSeq flowcell for paired-end 150 bp sequencing on the NovaSeq 6000 Sequencing System (Illumina) to obtain a mean read depth of *∼*100 million paired-end reads per sample. All samples were run twice, on both an S4 and S1 flowcell to achieve the targeted number of reads and minimise sequencing run bias. Sequenced reads were de-multiplexed and FASTQ files were generated using the BCL Convert software (Illumina). The pre-alignment quality control (QC) and alignment processes used in this study were previously described [48] and integrated into a Nextflow pipeline (https://github.com/Jbrenton191/RNAseq_splicing_pipeline). The data processing workflow used is summarised in **Supplementary Figure 2**.

#### Quality control and alignment

Fastp was used for adapter trimming as unprocessed sequencing reads in the FASTQ file may contain adapter sequences from oligonucleotide ligation to ends of cDNA fragments during the library preparation, and also used for read filtering and base correction through default settings^32^. Reads were filtered out if they had *≥*40 % Phred quality score < 15 (equating to a base call accuracy of 96.8 %) or with *≥*5 bases within a read that could not be called (‘N’). Reads shorter than 36 bases were removed. A base correction of a minimum 30 bp overlap was required with a total mismatch for fewer than 5 bases for the paired-end reads if one read in the pair had a high-quality score (Phred > 30) and the other a low-quality score (Phred < 15). FastQC (https://www.bioinformatics.babraham.ac.uk/projects/fastqc/)was used to visualise initial QC metrics of the fastp files before alignment. Reads from Run 1 (flowcell S4) and Run 2 (flowcell S4) showed consistent quality of sequencing between runs (**Figure 1**).

**Figure 1.**
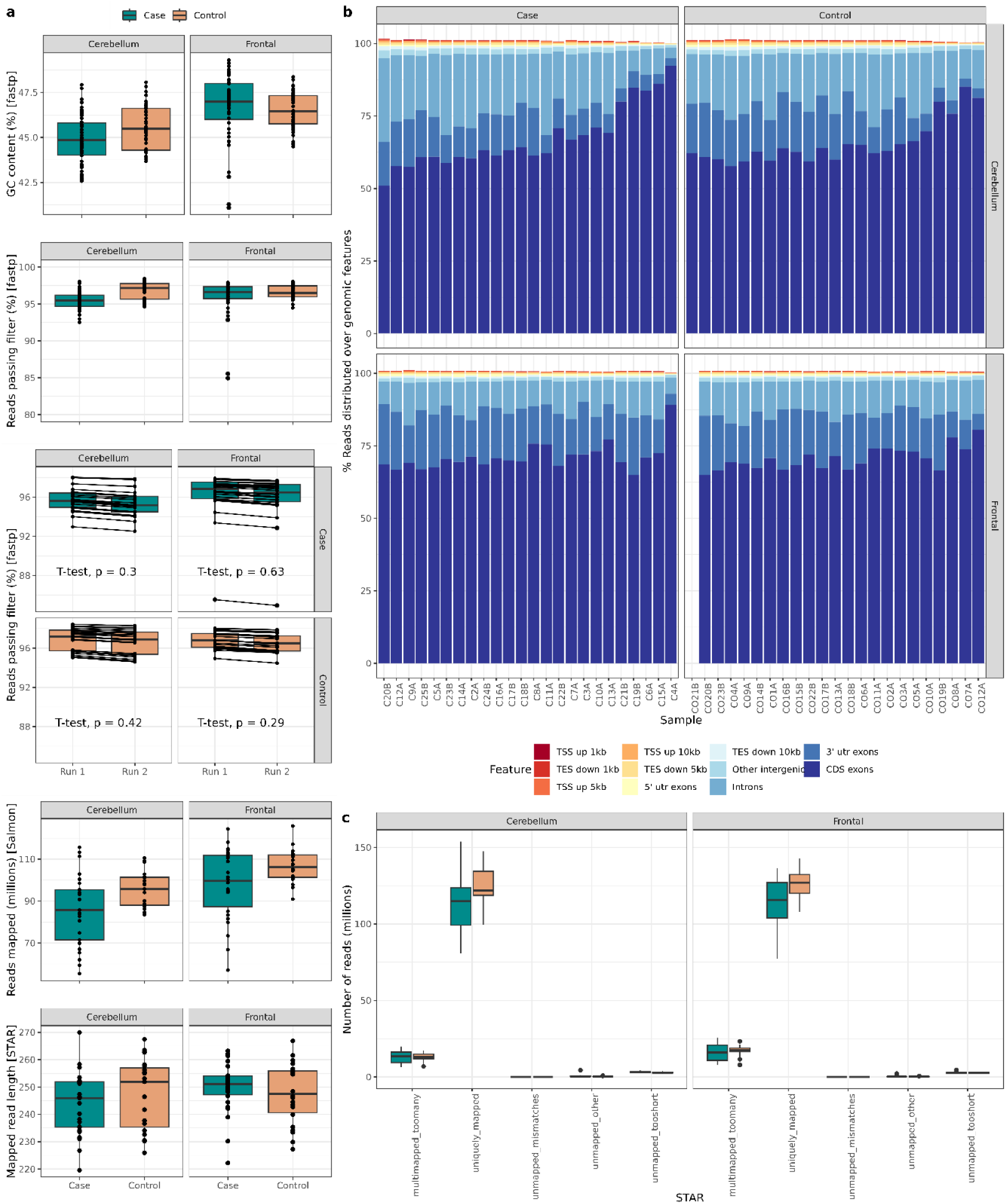
RNA-sequencing metrics. (**a**). RNA-sequencing metrics across ataxia cases and controls between the two brain regions from fastp (% GC content of reads, % reads passing filtering and comparison, % reads passing filtering between the two flowcell runs, with t-test p value comparing differences between the two runs), Salmon (number of millions of reads mapped per sample) and STAR (mean paired read length in base pairs, i.e. the paired-end read length is half of this value). For fastp results, reads from both runs are shown separately. Salmon and STAR results are derived from the merged fastp reads across the two runs. (**b**). Distribution of reads over genomic features across cases and controls, partitioned by brain region (cerebellum and frontal cortex). Genomic annotation: TSS represents the transcription start site and TES represents transcription end site. “Up” represents the sequence is mapped upstream of an annotation site such that “TSS up 1kb” represents annotation within 1 kilo-base pair upstream of the transcription start site. “Down” represents that the sequence is downstream of the site such that “TES down 10kb” represents annotation within 10 kilobases downstream of a transcription end site (**c**). Number of mapped reads following STAR alignment. Unmapped reads refer to reads that are not mapped as they are either too short, have a lot of mismatches or for another reason. Multimapped reads point to reads that map to multiple loci.

Spliced Transcripts Alignment to a Reference (STAR) software^33^ was used to map processed reads to the GRCh38 human reference genome, using gene annotations from Gencode v.38^34^. Two rounds of mapping were performed through individual 2-pass mapping to improve mapping of junction reads to novel splice junctions. Default settings were used for all STAR parameters, which adhered to standard ENCODE long-read RNA-sequencing guidelines (https://www.encodeproject.org/rna-seq/long-read-rna-seq/).

We used RSeQC quality metrics for post-alignment QC^35^ (**Figure 1**). A median of 240.7 million total paired reads per sample (range: 154.2 - 307.7 million reads), equating to a median of 120.35 million paired-end reads per sample was generated.

#### Quantification of bulk-tissue RNA-sequencing data

Salmon (v 0.14.1) was used to quantify processed reads with respect to the reference transcriptome, correcting for sequence-specific, fragment GC-content and positional biases (–seqBias, –gcBias, –posBias respectively)^36^. A *decoy-aware* transcriptome file based on GRCh38 and Gencode v.38 was generated using Alevin selective alignment (https://combine-lab.github.io/alevin-tutorial/2019/selective-alignment/). This avoids intronic reads being aligned to the transcriptome. The salmon index was created using the transcriptome fasta from gencode v.38 and the full genome used as a decoy (https://github.com/COMBINE-lab/salmon). *tximport*^37^ was then used to transform Salmon transcript-level abundance estimates to gene-level abundance estimates. Genes whose genomic co-ordinates were found to overlap ENCODE problematic regions^38^ were removed from downstream analyses to ensure adequate quality of the genes analysed. 41,636 genes were detected in cases and 42,192 genes in controls. Filtering removed genes with 0 counts in at least one sample across a disease or control group, resulting in 20,490 genes after filtering. Only genes detected post-filtering were used for ongoing quantification analyses. Following gene-level quantification, samples were checked for any mismatch between the reported sex of brain donors and the sex determined by the expression of sex-specific genes (*XIST* and *DDX3Y*). No mismatches were observed.

### Covariate selection

Principal components analysis (PCA) was applied to gene-level expression data to delineate sources of variation. Gene-level expression was applied to the 20,490 genes detected in all samples after filtering, that is, genes with a count > 0 across all samples, and transformed with variance stabilising transformation using DESeq2^39^. Pairwise correlations between experimental variables (Spearman’s for age at death and RIN; Kruskal Wallis for categorical variables: brain region, sex, and brain bank) and principal components (PCs) were calculated. Correlation p values were FDR-corrected (**Figure 2a**). Brain region was found to be most significantly associated with the first two PCs (**Figure 2b**). Subsequent analyses examined the two brain regions separately. Covariate correction was applied to the variables significantly associated with PCs. Samples were plotted by their first two PCs to determine how well disease groups separated before (**Figure 2c, d**) and after covariate correction. Following covariate correction for covariates showing significant correlation (FDR p < 0.05) with the PCs, namely RIN, PMI, sex, age of death and brain bank from which tissue was derived, there was a clear separation between ataxia cases (both diagnosed and unknown diagnosis) and controls within the cerebellar samples (**Figure 2e**). This separation was defined by the non-overlapping confidence ellipses indicating the probability of individuals belonging to a specified group with 95% confidence. For frontal cortex samples, there was distinction between molecularly diagnosed cases and controls (**Figure 2f**).

**Figure 2.**
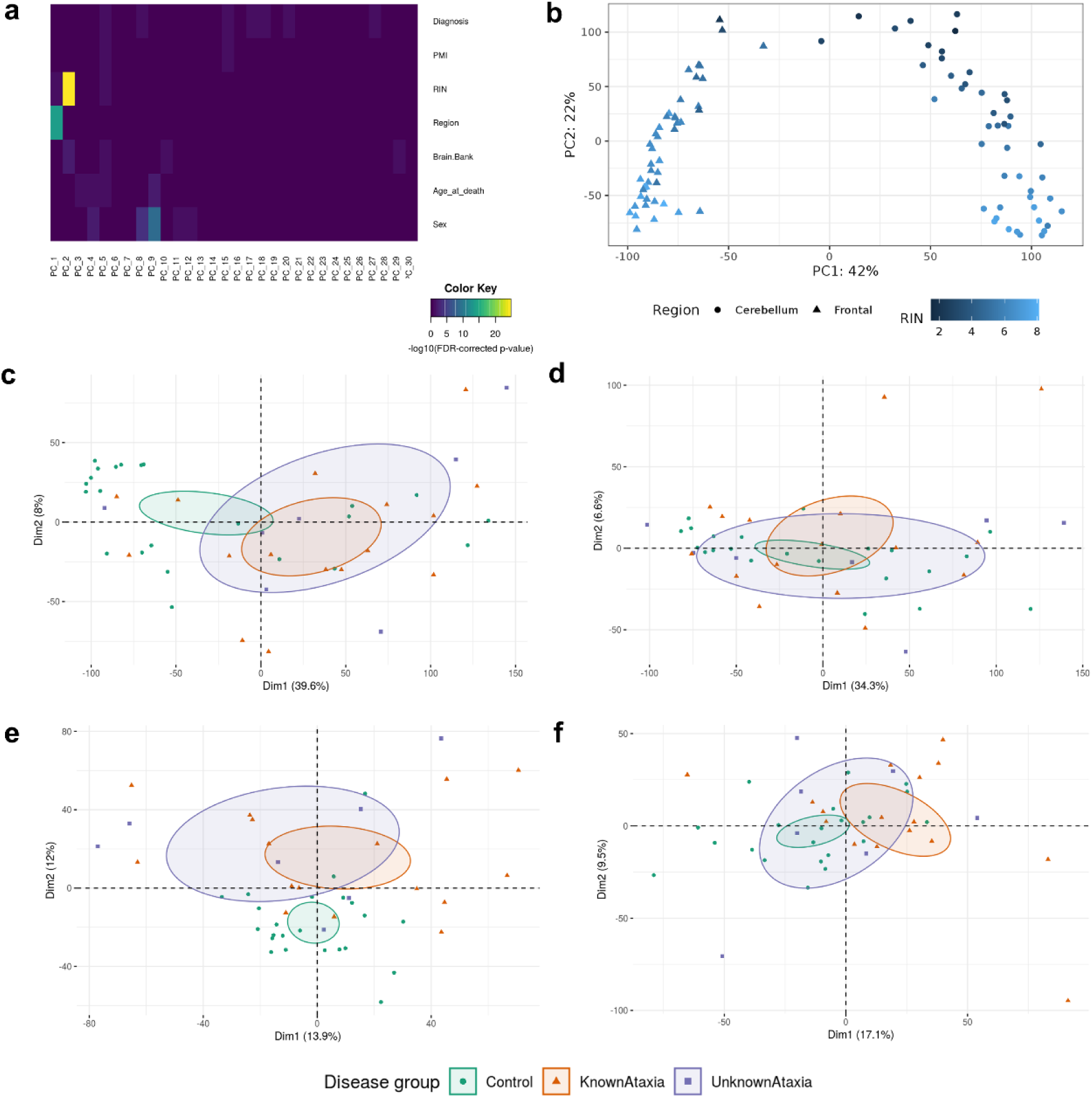
Principal components analysis (PCA) of gene-level expression. PCA was applied to the 20,490 genes detected in all samples after filtering. (**a**). The heatmap shows the significance of the correlations between sample numerical covariates and PCA results, and for categorical variables, the p value relating to the Kruskal Wallis test of the covariates with PCs. Significance was derived from Kruskal-Wallis for categorial variables and Spearman’s correlation for continuous variables. P values shown are FDR-corrected. The main principal components (PCs) of variation (PCs 1 and 2), explaining 42% and 22% of total variance respectively, were significantly correlated with brain region and RIN. (**b**). PCA plotting the first two PCs showing significant separation of brain region and RIN. (**c**). PCA of the first two PCs (Dim1 and Dim2) for cerebellar samples. The variance accounted for by each PC is in parentheses in the respective axes titles. The confidence ellipses indicate the probability of whether individuals belong in the specified group with 95% confidence. The groups are separated as controls, “KnownAtaxia” defined as those cases with known molecular diagnosis, and “UnknownAtaxia” defined as those cases without a known molecular diagnosis. (**d**). PCA of the first two PCs for frontal cortex samples. PCA in first two dimensions after covariate correction for RIN, PMI, sex, age-of-death and brain bank within the cerebellum (**e**). and frontal cortex (**f**), showing separation (based on the confidence ellipses) of all diagnosed (“known”) ataxia cases and controls.

### Overall framework for analysis

We applied three overall levels of analyses to gain additional insights through the different comparisons.

- Level 1: all cases were compared to all controls to gain an integrated understanding;
- Level 2: cases with a confirmed molecular diagnosis were compared to cases without a molecular diagnosis, and with controls, to provide diagnostic clues within those cases that have an unknown molecular diagnosis;
- Level 3: cases with specific genetic subtypes were analysed, namely SCA1 (n = 3), SCA2 (n = 3), SCA6 (n = 3) and FRDA (n = 4), and compared with controls.

Comparisons were made across the different levels across different diseases. Furthermore, comparisons were made at all the levels between the two brain regions: cerebellar and frontal cortices. Analyses focused on differential gene expression and differential splicing, partitioned through a transcriptome-wide approach and also targeted to the causative Mendelian gene of interest for those ataxias with a molecular diagnosis (**Figure 3**).

**Figure 3.**
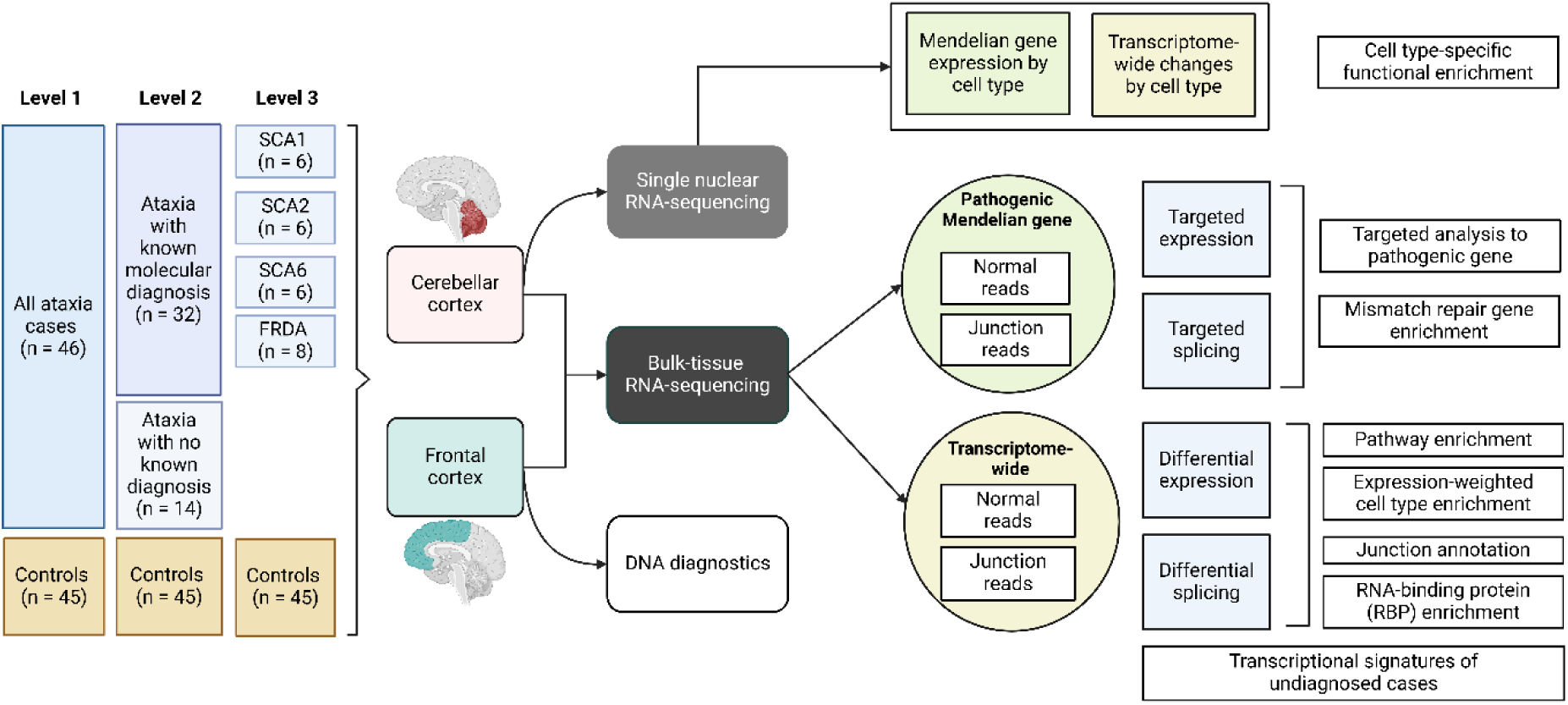
Overall framework for RNA-sequencing analysis. The 91 samples were partitioned into three levels of analyses: level 1 – cases compared to controls; level 2 – molecularly diagnosed cases compared with controls, molecularly undiagnosed cases compared to controls, molecularly diagnosed compared with molecularly undiagnosed cases; level 3 – cases with a molecular diagnosis where n *≥* 3 (SCA 1, 2, 6 and FRDA) compared with controls. Subsequent analyses focused more broadly either on quantification of the Mendelian gene of interest, or a through transcriptome-wide approach in analysing differential gene expression and differential splicing. Bul-tissue RNA-sequencing was carried out for both cerebellar and frontal cortices. Single nuclear (sn) RNA-sequencing was carried out for cerebellar tissues for Level 3 cases only. Image created in BioRender.

#### Bulk-tissue differential gene expression

Differential gene expression was assessed using the DESeq2 R package^39^ and for genes with count > 0 in all samples only (20,490 genes). Default parameters were used except for the maximum number of iterations allowed for convergence (maxit = 1,000). Gene-level expression was controlled for covariates selected after PCA. The covariate corrected model used the design formula:

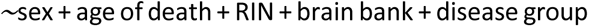

where disease group indicates the groups categorised at each of the analysis levels used for comparison. Therefore, the differential expression analysis was carried out for each of the levels, for example, comparing gene-level expression between cases and controls in Level 1, and comparing SCA1 and controls; SCA2 and controls; SCA6 and controls; FRDA and controls in Level 3 analyses. Pairwise comparisons were applied across the groups over the three levels of analysis to test gene-level differential expression with a log_2_fold change indicating the logarithmic fold change between the two conditions compared. A Wald test was used to test for significant differences between the two groups, such that the null hypothesis refers to no differences between the gene-level expression between the two groups. Multiple testing correction was performed by FDR-correction, with a significance cut-off < 0.05. The logarithmic fold change reported throughout represents the “shrunken” fold change reported in DESeq2. This allows more accurate fold change estimates when the gene count is low.

#### Bulk-tissue differential splicing analysis

Leafcutter was used to assess differential splicing^40^. Leafcutter detects reads with a gapped alignment (junction reads that span exon-exon junctions) that are representative of intron excision events, and uses them to quantify intron usage across samples without reliance on existing reference annotation or isoform estimation. In this way, Leafcutter does not rely on predefined transcript models such that potential pathogenic aberrant splicing events may be studied^40^. Local splicing events are constructed through intron clusters, where overlapping introns are connected by shared splice junction(s). STAR junction outputs (SJ.out.tab) were filtered to remove any regions that overlap ENCODE problematic regions^38^ and converted to the .junc files for intron clustering. Thresholds defining intron clusters were as follows through leafcutter_cluster.py:

– l 1,000,000A maximum intron length of 1,000,000 bps was applied as used in STAR.
– m 30introns detected in fewer than 30 reads across the samples.
– p 0.001fewer than 0.1% of reads in a cluster to support a junction.

Differentially spliced clusters were identified in a pairwise manner, controlling for the same covariates used in the design formula of differential expression listed above. Leafcutter uses a Dirichlet-multinomial generalised linear model to test for changes in the junction counts across introns within a cluster, and thus, differential intron usage across conditions. Clusters were annotated to genes using exon files generated from Gencode v.38. Within Level 1 and 2 analyses, only introns detected in *≥* 5 samples were tested and an intron cluster was only tested if detected in *≥* 3 individuals in each comparison group with an overall coverage of *≥* 20 junction reads. For Level 3 analysis, intron clusters had to be detected in *≥* 3 individuals in each comparison group to account for the smaller sample sizes per disease group. P values were FDR-adjusted. We defined a differentially spliced intron cluster as one with an FDR p < 0.05 that contained at least one intron with an absolute delta percent-spliced-in value (|ΔPSI|) *≥* 0.1. A total of 57,797 intron clusters were detected in 15,464 genes across all samples.

#### Annotation of introns by splicing event

Differentially spliced introns and introns within the Mendelian genes of interest were annotated using Gencode v.38 using junction_annot() from dasper package^41^. This function categorises junction reads based on the refGenome package annotation (https://github.com/cran/refGenome). Junctions were accordingly classified as: annotated; novel exon skip; novel combination; novel acceptor; novel donor; ambiguous gene and unannotated. Annotated junctions have both donor and acceptor splice sites that match the boundaries of an existing intron. Novel exon skip and novel combination junctions have donor and acceptor splice sites that overlap known exon boundaries but not of constitutive exons within the same transcript or different transcripts respectively. Novel donors (3’ end) and novel acceptors (5’ end) are junctions where only one end matches an existing exon boundary, so are partially annotated to the reference transcriptome. Unannotated junctions have no overlaps with known exon boundaries (**Supplementary Figure 3**). Junctions that mapped to more than one gene (“ambiguous gene”) were not considered due to ambiguity of interpretation. Splicing events within Mendelian genes of interest were visualised using the ggtranscript package^42^ after junctions were directly extracted from the BAM files. Differentially spliced introns within the Mendelian genes of interest were also visualised using the same package.

#### Expression and splicing of associated Mendelian genes

For cases of ataxia with known molecular diagnosis, we specifically reviewed the causative Mendelian genes of interest for expression, namely: *ATXN1* (SCA1); *ATXN2* (SCA2); *CACNA1A* (SCA6); *ATXN7* (SCA7); *TBP* (SCA17) and *FXN* (FRDA). First, for differential expression of these genes, the DESeq2 output comparing the targeted gene-level expression between cases and controls was used. For direct quantification of gene expression, normalised coverage of the Mendelian gene for a particular sample was calculated as:

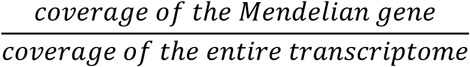

The coverage was directly extracted from the aligned BAM files using the GenomicRanges package^43^ and defined as the number of counts in the sequencing data overlapping particular genomic positions. Genomic co-ordinates of the Mendelian gene of interest were extracted from Gencode v.38. For exon-level coverage, termed normalised exon coverage, the coverage of the exon (Gencode v.38-defined genomic coordinates) was normalised for the number of base pairs across that exon. Lastly, coverage was also estimated for the repeat-containing region of these genes with coordinates determined by PanelApp (https://panelapp.genomicsengland.co.uk/)^44^. For splicing events, the junction reads were quantified for the annotated splicing event by reviewing the sum of read counts for a particular junction. To assess whether genes known to be associated with ataxia are enriched within differentially expressed and differentially spliced genes, a chi-squared test was performed and an FDR p < 0.05 deemed to be significant.

#### Functional enrichment analyses

Gprofiler2 for gene set enrichment of Gene Ontology (GO) biological process, molecular function and cellular component, Reactome and KEGG terms was used^45^. For differentially expressed genes, the background gene set comprised all 20,490 genes tested for differential expression. For differentially spliced genes, the background gene set comprised 15,464 unique genes in which intron clusters were identified in all samples. Functional enrichment analyses were performed at all three levels for differentially expressed and differentially spliced genes, and across each of the brain regions. P values were adjusted using the g:SCS correction method, accounting for overlap in functional terms, with a significantly enriched term having a multiple-testing corrected p value < 0.05. Similarities and differences in the number of enriched terms were visualised using the UpSetR package^46^. Terms were also summarised semantically using function go_reduce (https://github.com/RHReynolds/rutils/blob/master/R/go_reduce).

#### Expression-weighted cell type enrichment

Expression-weighted cell type enrichment (EWCE) assesses whether the expression of a particular set of genes is enriched within a particular cell type than would be expected by chance^47^. EWCE was used to assess whether sets of differentially expressed or differentially spliced genes were more likely to be expressed within a certain cell type. The gene cell type specificity values derived from single-cell or single-nuclear RNA-sequencing data are contained within a specificity matrix and required as input data for EWCE in addition to the tested gene sets. Two different specificity matrices were used: a human post-mortem brain single-nuclear RNA-sequencing dataset for cortical cell types (https://portal.brain-map.org/atlases-and-data/rnaseq/human-mtg-smart-seq) and a mouse single-cell RNA-sequencing dataset which contains multiple brain regions, including cerebellar-specific cell types^48^. Further descriptions of the cell types used in the former dataset are available in **Supplementary Table 3**. For each combination of gene list and specificity matrix, 10,000 bootstrap replicates were used. Transcript length and GC-content biases were controlled for by using bootstrap replicates with similar properties as the tested gene lists. The output generates a standard deviation from the mean, and a logarithmic fold change, which indicates the difference between the target gene list’s mean expression and bootstrap replicates’ mean expression, and the fold change between the expression of the target gene list and bootstrap replicates respectively. FDR multiple testing correction was applied to account for the total number of cell types tested within each specificity matrix and the number of comparisons made.

#### RNA-binding protein differential expression and splicing

As RNA binding proteins (RBPs) play important roles in the pathogenesis of repeat expansion disorders, we studied the differential expression and differential splicing of RBPs in ataxia patients compared with controls. We used the EMBL RBP database v.0.2.1 (https://rbpbase.shiny.embl.de/)^49^ where a superset of 3,470 RBPs are defined with information from previously annotated RBPs, RNA interactome capture studies (RIC) and mass spectrometry data. This superset consists of 2,650 RBPs that have been detected in at least two RIC studies. We assessed the differentially expressed and spliced genes for RBP function using these predefined databases.

### Single nuclear RNA preparation and sequencing

#### Isolation of nuclei, nuclei encapsulation and snRNA-sequencing data generation

Cerebellar samples were used for single nuclear RNA-sequencing (snRNAseq). Before starting the nuclei extractions, ultracentrifuge SW 41 Ti Swinging-Bucket Rotors (Beckman Coulter) were cooled down in the Optima XPN-90 Ultracentrifuge (Beckman Coulter) at 4 °C to prevent degradation of RNA. Buffers were prepared from stock according to this protocol (https://www.protocols.io/view/nuclear-isolation-of-post-mortem-brain-tissue-for-yxmvm25xng3p/v1) and kept on ice until its use. Digitonin from this protocol. Homogenisation involved using 2mL Tissue Grinder Homogenisation Glass Vessel (VWR International) and 2mL Tissue Grinder Plunger (VWR International) with ice-cold HB (Homogenisation Buffer). Tissue was homogenised with 80 gentle strokes. The homogenate was then transferred to an ice-cold falcon tube, with additional 650uL HB added, followed by adding 2.65mL Gradient Medium (GM). The mixture was gently mixed by flipping, without vortexing. Ultracentrifuge tubes (Beckman Coulter) were first layered with 4mL of 29% Cushion buffer, followed by adding 5mL of the sample on top. The samples were balanced by weight and then spun in an ultracentrifuge at 7700 RPM at 4 °C for 30 minutes.

After centrifugation, the supernatant was carefully removed without disturbing the nuclei pellet using a Pasteur pipette. The nuclei were then resuspended in RNaseq Wash Buffer, pooled, and filtered through 5 ml Round Bottom Polystyrene Test Tubes with Cell Strainer (SLS). The sample was then centrifuged with the table-top centrifuge (Eppendorf) at 500 RCF, 4 °C for 5 minutes. The supernatant was removed, the nuclei were resuspended again and spun down with the same conditions in table-top centrifuge. Finally, supernatant was removed and nuclei were resuspended in 400 µl of RNaseq Wash Buffer. Nuclei were counted using C-Chip Neubauer Hemocytometer (Cambridge Bioscience) and light microscope, as well as LUNA-FL cell counter (Logos Biosystems) with Acridine Orange (Invitrogen) staining.

All samples were processed as per Chromium Next GEM Single Cell 3ʹ Reagent Kits v3.1 (Dual Index). Following manufacturer’s guidelines, the samples were processed to target 7,000 nuclei per sample. We performed 11 cycles of cDNA amplification and 14-16 cycles of final indexing PCR, depending on the available cDNA concentrations. The concentrations and quality of cDNA and indexed final libraries were measured using D5000 ScreenTape Assay and D1000 ScreenTape Assay, respectively, on 4200 TapeStation (Agilent) instrument. The samples were sequenced on Novaseq X Plus on 25B or 10B flow cells with a target of 100 bp paired-end reads.

#### Single nuclear RNA-sequencing data processing and cell type identification

Cell Ranger^50^ was used to convert raw BCL sequencing files to fastq files and align raw sequencing reads to the GRCh38 (ENSEMBL v.107) human reference genome. QC, pre-processing, batch correction and clustering of nuclei were performed using the Panpipes package^51^. Doublet content was estimated using Scrublet^52^, as well as estimating gene expression metrics, namely mitochondrial and ribosomal RNA content. These outputs were used for selecting filtration parameters, leaving a total of 133,000 high quality nuclei. Seurat^53^ was used to identify highly variable genes and batch correction was performed using Harmony^54^ and community detection with the Leiden algorithm. Major cell types were annotated based on marker expression across the clusters, summarised in **Supplementary Table 4**. The final number of nuclei and genes analysed per cell type is summarised in **Supplementary Table 5**. The remaining nuclei were visualised using a non-linear dimensionality reduction algorithm Uniform Manifold Approximation and Projection (UMAP, v.0.1.10).

Differential gene expression was analysed in this dataset using the Dreamlet R package^55^. This package applies linear mixed modelling to pseudobulked data, denoting raw count matrices that have been summed for a given sample and cell type. The voom package^56^ then normalises pseudobulked counts per cell type, setting the minimum number of reads for a gene to be considered expressed to 10. Differential gene expression analysis is then performed using the limma^57^-style set up of Dreamlet’s predecessor, Dream^58^. Contrasts were made between controls and each diagnosis group, as well as all four ataxias grouped together (**Supplementary Table 6**). Multiple test correction was performed using FDR correction, with a significance cut-off of < 0.05.

## Results

### Clinical phenotyping and data structure

To study commonalities across the hereditary ataxias, we analysed brain samples from a total of 23 individuals with a clinical diagnosis of ataxia and 23 controls. Of those with a diagnosis of ataxia, sixteen cases had confirmed molecular diagnoses consistent with those recorded by the brain banks and were as follows: Friedreich’s ataxia (n = 4); SCA1 (n = 3); SCA2 (n = 3); SCA6 (n = 3); SCA7 (n = 1); SCA17 (n = 2). The remaining seven cases did not have expansions in any of the common repeat disorders (SCA1, 2, 3, 6, 7, 17, NIID, CANVAS, SCA27b) and had no pathogenic variants detected on whole exome sequencing, and were categorised under ataxia without molecular diagnosis (“unknown cases”). PCR-based estimates of the repeat expansion sizes for the SCAs across all samples are shown in **Supplementary Figure 4**. Paired frontal and cerebellar cortex tissue were retrieved from each donor (except in one case where only cerebellar tissue was available), resulting in a total of 46 cerebellar and 45 frontal cortex tissue samples. Age and sex were matched between cases and controls (**Supplementary Table 7**). The variable ages at death between different types of ataxia (Kruskal Wallis p = 3.3 *×* 10*^−^*^6^) (**Supplementary Figure 4**) reflect the clinical heterogeneity of disease. Individuals with SCA6 have the oldest age at death, consistent with the later onset and slower progression of disease^59^.

### Widespread differences in the transcriptional signatures of disease across brain regions

Given the phenotypic and genetic similarities across the hereditary ataxias with the majority caused by pathogenic repeat expansions, we postulated that this would result in common transcriptomic signatures that would be most evident in the cerebellum. To explicitly test this hypothesis, we analysed transcriptomic data generated across all samples at the three levels of analysis. Transcriptome-wide differential expression analyses identified distinct signatures across the different levels of comparison and disease subtypes (**Figure 4**).

**Figure 4.**
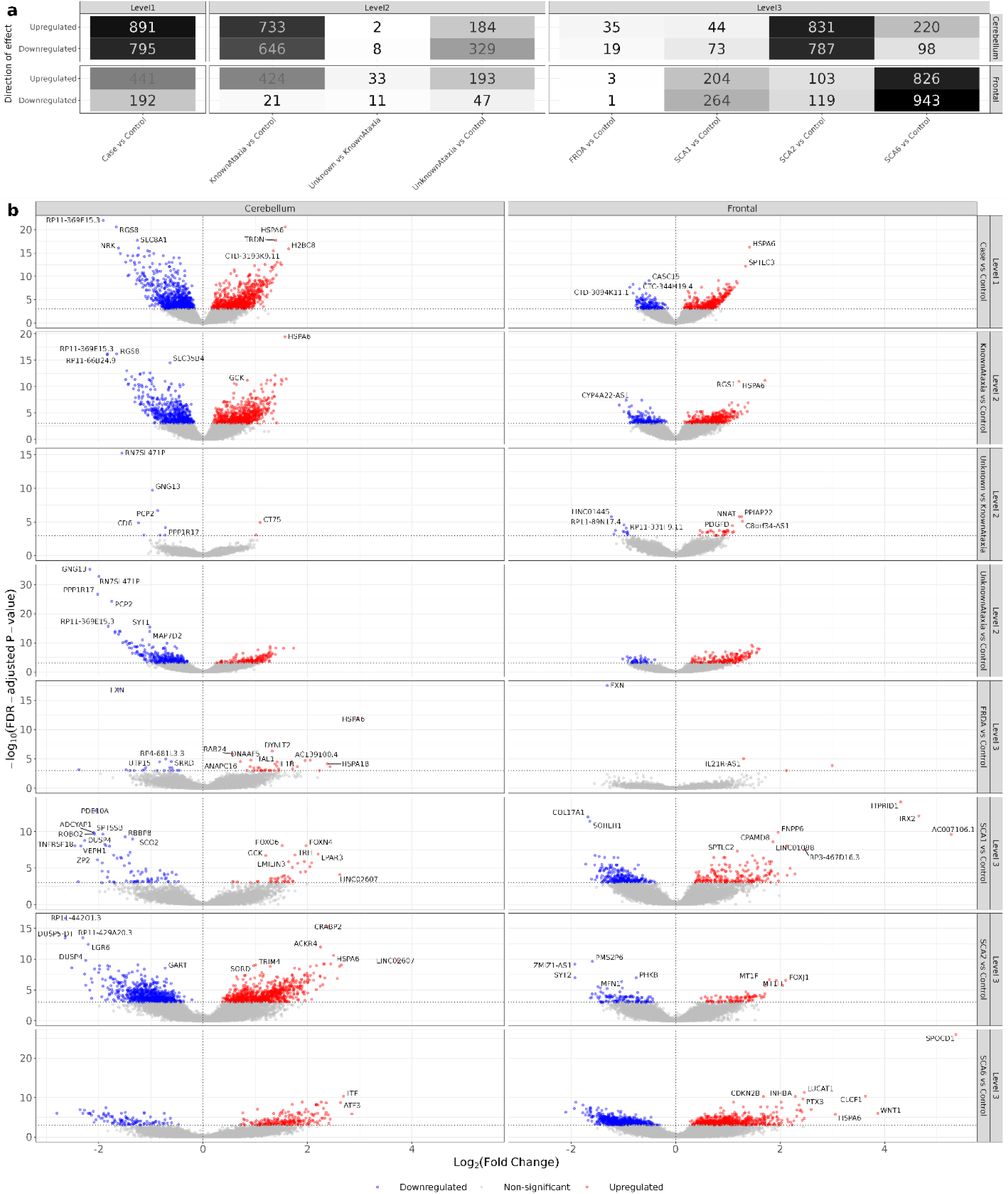
Differential gene expression. Differentially expressed genes in all disease comparison groups across two brain regions. **(a).** Total number of differentially expressed genes (FDR p < 0.05) across the comparison groups in cerebellar and frontal cortices. Darker shading corresponds to a higher number of differentially expressed genes. **(b)**. Differential expression of genes across the individual comparison groups. Genes with the highest fold changes within each comparison set are annotated with the gene symbol. The vertical dashed line shows that there is no change in expression between the two comparator groups and the horizontal line represents an equivalent FDR p of 0.05. The three levels of analysis relate to: Level 1 – cases compared to controls; Level 2 – molecularly diagnosed cases (“KnownAtaxia”) compared with controls, molecularly undiagnosed cases (“UnknownAtaxia”) compared to controls, molecularly diagnosed compared with molecularly undiagnosed cases; Level 3 – cases with a molecular diagnosis where n *≥* 3, compared with controls (SCA 1, 2, 6 and FRDA).

First, there were more differentially expressed genes within the cerebellum compared to frontal cortex across most comparisons (**Figure 4a**), consistent with regional differences in pathology. However, this was much less pronounced than expected with unexpectedly high numbers of both up- and downregulated differentially expressed genes within the frontal cortex. In particular, many differentially expressed genes were found in SCA6 frontal cortex compared to controls (**Figure 4a**) with large effect size (**Figure 4b**, **Figure 5**). As SCA6 is a clinically pure cerebellar syndrome^59^, the extent of the transcriptomic response in frontal cortex was particularly unexpected, and potentially suggested that we were capturing compensatory processes.

**Figure 5.**
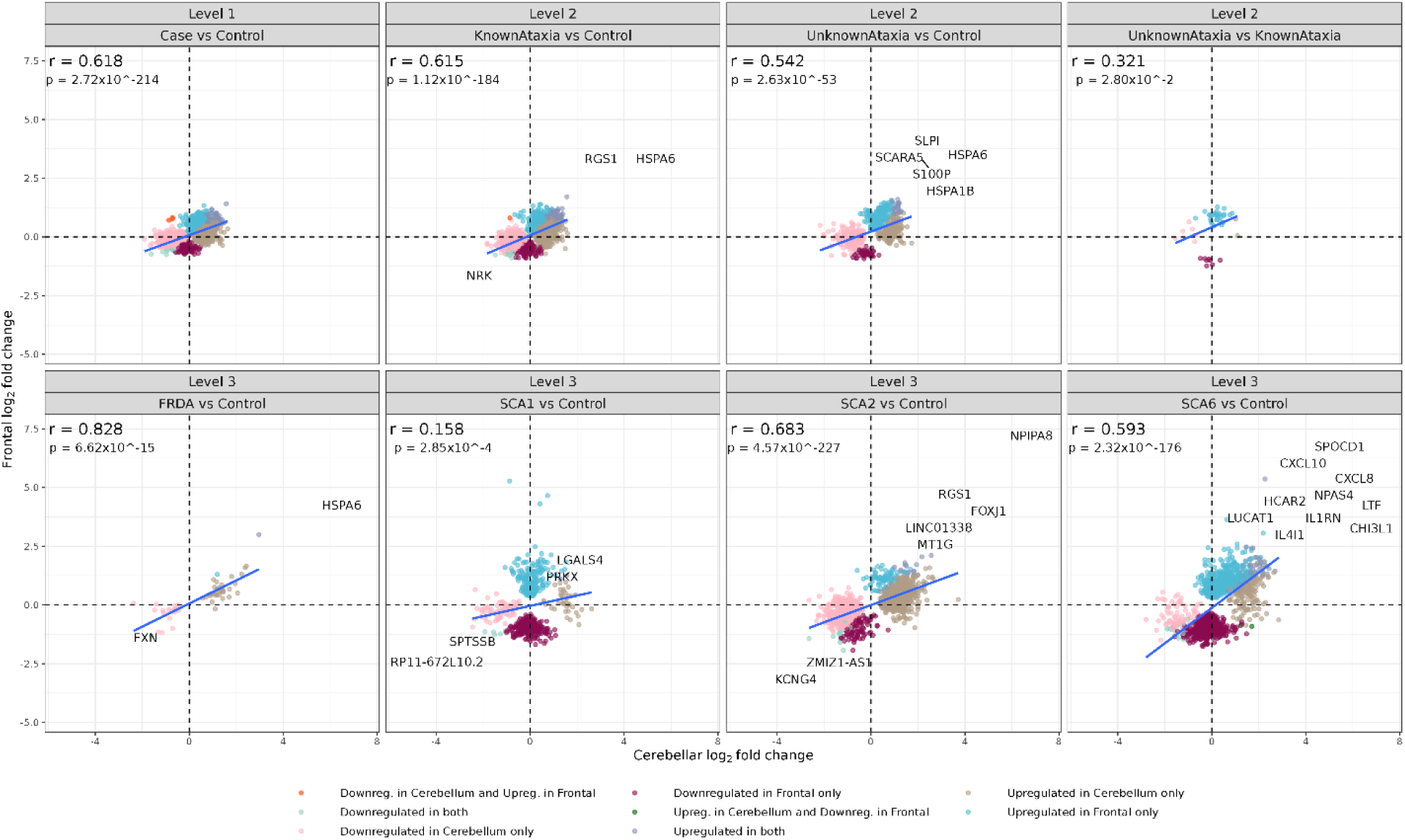
Cerebellar-frontal expression correlation. Comparison of the fold change between all significantly differentially expressed genes (FDR p < 0.05) within the cerebellum and frontal cortex across the disease comparisons. Genes upregulated or downregulated in both regions and have the highest fold change are annotated. The *r* for each plot represents the Pearson’s correlation coefficient for cerebellar-frontal fold change in gene expression for that particular disease comparison, and p is the corresponding p value.

In fact, this led us to our second observation, namely that transcriptomic responses across both brain regions and all levels of analysis were characterised not only by the downregulation of gene expression asmightbe expected in neurodegenerative diseases, such as hereditary ataxia, but also upregulation of gene expression. The latter may indicate the activation of processes to mitigate disease effects (**Figure 4**).

Thirdly, there were distinct transcriptional signatures in molecularly unknown ataxia cases compared to controls (**Figure 4**). The similarity in gene expression between molecularly unknown and known cases was most apparent in the cerebellum. While the similarity in downregulated transcriptional signatures in the cerebellum is expected, the similarity in upregulated genes within the cerebellum is less obvious suggesting that both molecularly known and unknown cases may have similar compensatory mechanisms. Lastly, there were very few differentially expressed genes within FRDA compared with controls in both brain regions. The number of gene expression changes within the other Mendelian ataxia disorders suggest more widespread transcriptional changes and involvement in RNA processing.

### High correlations in transcriptomic signatures across affected and relatively unaffected brain regions

In order to ascertain the relationship between gene expression within the cerebellum and frontal cortex, we compared gene expression fold change estimates of both regions across each of the disease comparisons (**Figure 5**). Overall, there was a statistically significant positive correlation (Pearson’s correlation coefficient, *r*, corresponding p value in **Figure 5**) between cerebellar and frontal cortex change in expression (*r* range between 0.158 and 0.828). The weakest correlation was found in SCA1 compared to controls (*r* = 0.158, p = 2.85 × 10*^−^*^14^) indicating that some genes were only up- or down-regulated within one region. Interestingly, the cerebellar-frontal fold change correlation was high (*r* = 0.593, p = 2.3 × 10*^−^*^176^) when comparing gene expression in SCA6 to controls. This is despite SCA6 being considered clinically to be a “pure” ataxia syndrome, with disease presumed to be restricted to the cerebellum.

### Pathway analysis implicates the immune system in disease

Functional enrichment analyses found varying numbers of significantly enriched (defined as gSCS-corrected p < 0.05) pathways at each level of comparison (**Figure 6a**). Of interest, there were numerous enriched pathways amongst upregulated genes in the frontal cortex in ataxia cases. These upregulated pathways were most apparent within SCA6 frontal cortex compared with controls.

**Figure 6.**
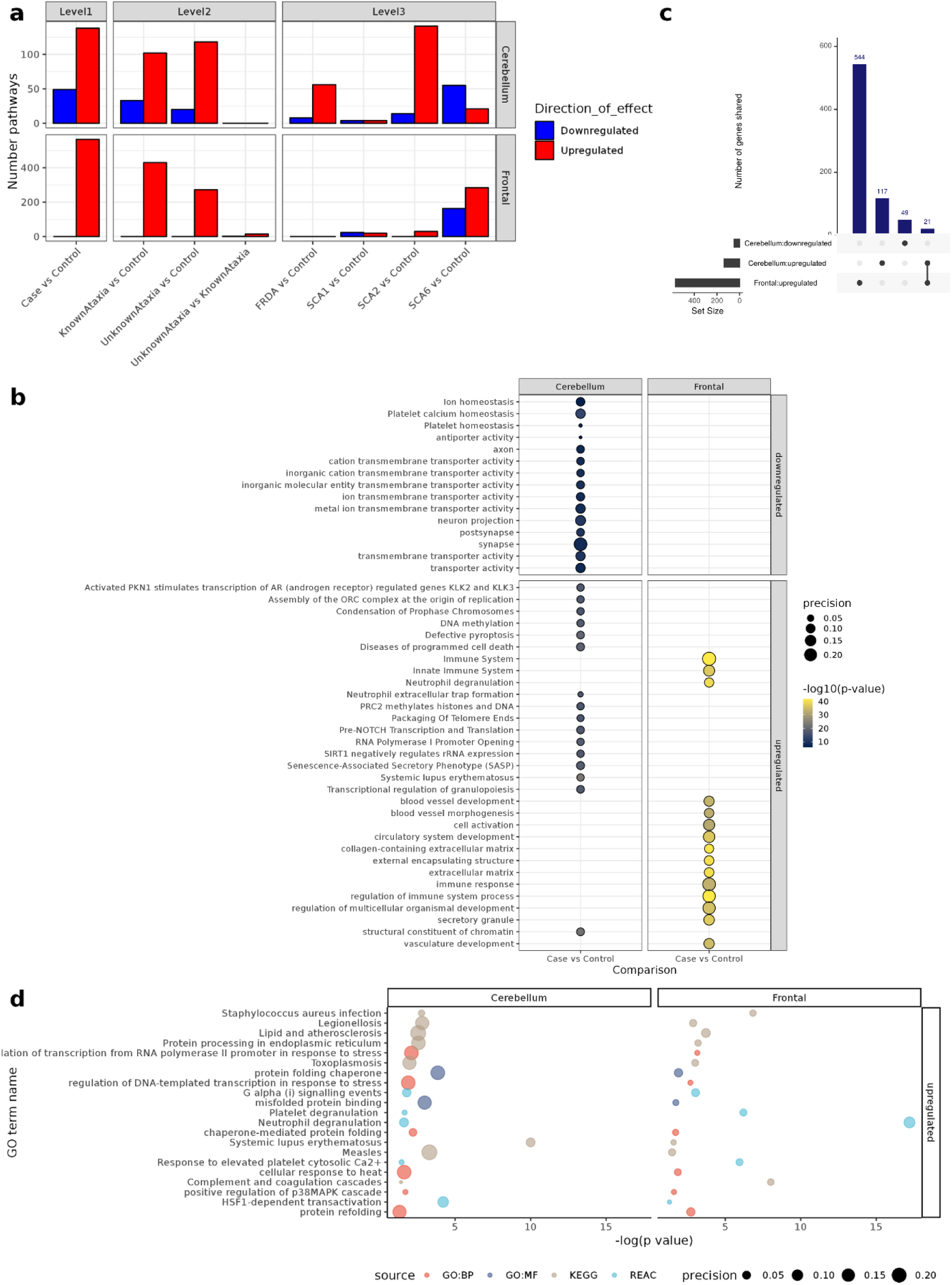
Pathway enrichment of differentially expressed genes. (**a**). Total number of significantly enriched pathways across disease groups. (**b**). Top significantly enriched gene ontology (GO) terms in cases compared to controls aross two brain regions. (**c**). Similarities and differences in enriched pathways between different brain regions in cases compared to controls. (**d**) Overlapping GO terms between two brain regions in cases compared to controls. Source refers to the pathway annotation category. Precision is from gprofiler package and defines the proportion of genes in the input gene list that can be assigned this term. P values given are gSCS-corrected.

Focusing on pathways associated with differentially expressed genes in all cases compared to controls, downregulated pathways within the cerebellum were associated with synaptic and transporter activity, suggesting dysfunction within these pathways (**Figure 6b**). The most significantly upregulated pathways within the cerebellum of ataxia cases compared to controls were related to DNA structure (chromatin, packaging of telomeres, histone methylation) or immune-mediated processes involving white blood cells such as neutrophil degranulation and transcriptional regulation of granulopoeisis. Of interest, alcohol biosynthetic and metabolism pathways were also significantly upregulated in the cerebellum of cases compared to controls. The upregulation of methylation pathways included the polycomb repressive complex 2 (PRC2) methylation pathway, that has been shown to silence genes associated with neurodegeneration [78]. Upregulated pathways within the frontal cortex related both to the innate immune system and its regulation, as well as vascular development. Of particular note, there was an enrichment of interleukin (IL) pathways including IL-4, IL-6, IL-8, IL-10, IL-13, and IL-17. In line with this, only 21 terms overlapped between upregulated cerebellar and frontal cortex pathways (**Figure 6c**) and were mostly related to immune function and protein folding (**Figure 6d**). This suggests the importance of immune system activation in ataxia.

There was high similarity across down- and upregulated cerebellar pathways across molecularly undiagnosed and diagnosed ataxia cases (**Supplementary Figure 5a**). Molecularly undiagnosed cases also had distinct and unique pathways when compared to diagnosed cases within the upregulated pathways of the frontal cortex. Neurodevelopmental terms were significantly upregulated within undiagnosed cases, shedding light on the underlying biology of disease.

Furthermore, although there was distinct pathway enrichment between different genetic subtypes of ataxia, they all had upregulated immune pathways (**Supplementary Figure 5b**). Interestingly, pathways relating to innate immune responses were partially upregulated in the frontal cortex of SCA1 individuals and appear to be driven by genes such as *CPAMD8*, a member of the complement 3/alpha(2) macroglobulin family, with important roles in innate and adaptive immunity^60^. Within FRDA, there was a downregulation of heme processing within the cerebellum consistent with the putative function of the associated *FXN* gene. There was also an upregulation of IL8, myeloid cell differentiation and alcohol metabolism-related pathways within the cerebellum. In addition, pathways associated with legionellosis, measles, and toxoplasmosis were enriched.

In SCA2 compared to controls, there was downregulation of ion transporter activity within the cerebellum. Upregulated pathways in the cerebellum include protein folding, alcohol metabolism, DNA methylation including PRC2 pathways, gene silencing by RNA, RNA polymerase and transcriptional regulation. Of interest, immune signatures as reflected by legionellosis, measles, toxoplasmosis and granulopoiesis pathways were also highlighted. Furthermore, the enrichment of upregulated base excision repair, DNA damage and recruitment of ATM-mediated phosphorylation pathways provide insight into the mechanisms of disease within the SCA2 cerebellum. Given that DNA mismatch repair pathways are implicated as a disease modifiers in CAG-repeat expansion diseases through modulating somatic repeat instability, we investigated whether there was evidence of differential expression of mismatch repair and base excision repair genes as previously defined. We found evidence of differential expression of these genes across different brain regions and ataxia subtypes (**Supplementary Figure 6**). Notably, *MSH3* was significantly downregulated in the ataxia cerebellum but not frontal cortex. Likewise, in the SCA2 frontal cortex, there was upregulation of response to ions, regulation of the immune response and alcoholic liver disease.

SCA6 is associated with CAG repeat expansion in *CACNA1A* and is associated with downregulation of calcium signalling pathways within the cerebellum and upregulation of immune system pathways in both the cerebellum and frontal cortex. However, in the frontal cortex, the immune system upregulation mainly involved the innate immune system, namely cytokine activation, leukocyte regulation and interleukin activation pathways. Within the SCA6 cerebellum, there was an upregulation of both the adaptive and innate immune response, with the latter incorporating cytokine regulation and the complement cascade. *SPOCD1* was upregulated in both the frontal and cerebellar cortex of SCA6 patients compared to controls. The log fold change in expression of *SPOCD1* was higher in the frontal cortex (log_2_fold change = 5.36, FDR p = 5.01 × 10*^−^*^12^) compared with the cerebellum (log_2_fold change = 2.26, FDR p = 0.00014) (**Figure 5**). SPOCD1 is a nuclear protein whose expression is thought to be restricted to the period of *de novo* genome methylation, and is essential for piRNA-directed methylation and silencing of young transposable elements including long interspersed nuclear element-1 (LINE1)^61^.

The upregulation of alcohol metabolism-related pathways in all ataxia studied here except SCA6 is of interest as alcohol misuse is associated with cerebellar dysfunction suggesting an overlap in mechanism with hereditary ataxia in the underlying biological pathways.

In summary, there is a collective and unexpected upregulation of immune-associated pathways in both the cerebellum and frontal cortex. Downregulation of genes associated with transporter dysfunction and synaptic pathways is present only within the cerebellum, and not within the frontal cortex, showing cerebellar dysfunction in disease.

### Differential expression of causative Mendelian genes

As most cases of molecularly diagnosed hereditary ataxia are explained by monogenic variants, we investigated any association between expression of causative Mendelian genes and brains of donors with ataxia recruited in this study. To do this, I took two approaches: first, I reviewed the differential expression of the causative Mendelian genes in cases of ataxia compared to controls; secondly, I extracted the normalised coverage of the causative gene. More explicitly, we wanted to look for any loss of function as a potential pathogenic mechanism

Pathogenic expansions in *FXN, ATXN1, ATXN2, ATXN7, CACNA1A* and *TBP* are associated with FRDA, SCA1, SCA2, SCA7, SCA6 and SCA17 respectively. Taking the genomic coordinates to define the location of these repeat expansions, I found that there was no read coverage in the repeat location in either control or ataxia brains likely due to multimapping of these repetitive sequences. However, the gene-level and exon-level coverage (where the exon of interest is that in which the repeat is located) was not affected by the presence of a repeat expansion.

The only significantly differentially expressed causative gene within cases of the same subtype was *FXN*, which was downregulated in FRDA patients compared to controls (**Figure 7a**). This was supported by analysis through pairwise comparisons of the normalised gene coverage of each of the disease groups with controls, with a lower normalised coverage of *FXN* in FRDA samples compared to controls in the frontal cortex (Wilcoxon p = 0.0014, p = 0.058 for cerebellum samples). Reduced blood and fibroblast *FXN* mRNA levels have been reported in FRDA patients, with inverse correlation to GAA repeat size^62–64^. Resulting low frataxin protein has been described in the cerebellum and negligible levels in the cortex of individuals with FRDA^65^. However, CNS *FXN* expression is poorly described to date. The downregulation of *FXN* in both cerebellar and frontal cortices here are consistent with current understanding of *FXN* downregulation secondary to epigenetic silencing^66^. The lack of differential expression of the causative genes of the other Mendelian disorders suggests that for the other repeat expansion disorders, gene expression is not a critical component of pathogenesis, and loss of function is unlikely to be the dominant pathogenic mechanism. Furthermore, the upregulation of *ATXN1* in SCA2 cerebellum and downregulation of *CACNA1A* in SCA1 frontal cortex support their involvement in the same pathways and functional networks in the pathogenesis of ataxia, or the association of genes known to cause ataxia with cerebellar-specific function.

**Figure 7.**
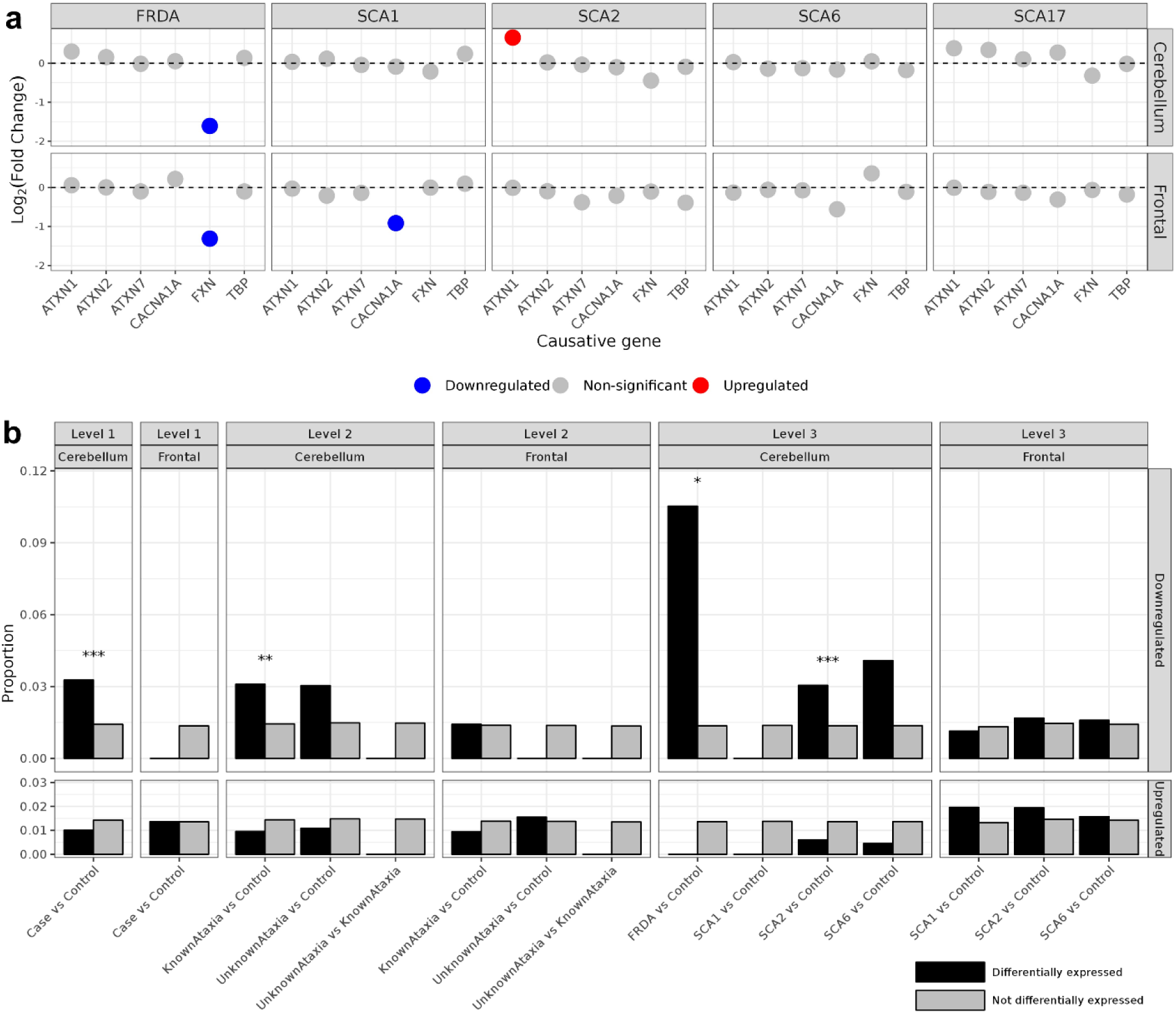
Differential expression of causative Mendelian genes and proportion of known ataxia genes. (a). Differential expression of causative Mendelian genes associated with molecularly diagnosed cases of ataxia in this study. **(b).** Comparing the proportion of known ataxia-causing genes defined in [5] between significantly differentially expressed genes (FDR p < 0.05) and genes not significantly differentially expressed within the comparator group. Comparisons between the proportions were made using chi-squared test and significant results are indicated as follows: ***FDR p < 0.001; **FDR p < 0.01; *FDR p < 0.05.

Hereditary ataxias are one of the most genetically heterogeneous disorders. Taking the 318 genes in which pathogenic variants are known to be associated with ataxia compiled from a combination of PanelApp, GeneReviews and OMIM^67^, we investigated the intersection between these known ataxia-associated genes and differentially expressed genes from bulk RNA-sequencing data (**Figure 7b**). We found significant (FDR p < 0.05) enrichment in known ataxia genes amongst genes downregulated within the cerebellum of individuals with ataxia compared to controls (**Figure 7b**). This downregulation was specific to FRDA and SCA2. These findings support that either downregulated cerebellar genes have specific cerebellar neuronal expression or that these genes may play a role in an co-expression network. There were no significant differences in the proportion of known ataxia genes within differentially expressed genes of the frontal cortex or upregulated genes of the cerebellar cortex.

### Upregulation of glial cell type-specific expression in ataxia brain

In order to assess the significance of differentially expressed genes at the cell type level, EWCE was used to ascertain which genes in an input list were more likely to be expressed by a particular cell type than expected by chance. In general, downregulated genes were enriched for neuronal cell type-specific expression in both the cerebellum and frontal cortex (**Supplementary Figure 7**). An upregulation of genes with glial cell type-specific expression was seen in both the cerebellum and frontal cortex, namely, in microglia, astrocytes, and oligodendrocytes (**Supplementary Figure 7a**). Interestingly, there was an enrichment of genes with microglial-specific expression in the frontal cortex of all ataxia cases. In SCA1 frontal cortex, there was particular enrichment of oligodendrocyte cell type-specific expression over other glial cells. In SCA2, microglial-specific expression was seen, and in SCA6 frontal cortex, all glial cells had prominent cell type-specific expression. The upregulation of genes with microglial-specific expression was consistent with an upregulation of genes associated with immune and inflammatory pathways. As the cerebellum contains unique cell types not be captured in other brain regions, the analysis was extended to focus on cerebellar neuronal cell types. We found downregulation of genes with specific expression across all cerebellar-specific neuronal cell types (**Supplementary Figure 7b**). More specifically, genes with Purkinje cell type-specific expression were significantly enriched within cerebellar downregulated genes, reflecting the importance of Purkinje cells in disease.

### Widespread differential splicing in the disease state

Using the EMBL RBPbase that catalogues a high number of genes associated with RBP function, I found evidence for differential RBP gene expression across different disease subtypes and brain regions (**Supplementary Figure 8**). Of note, downregulation of genes associated with RBPs was most evident within the cerebellum rather than frontal cortex (**Supplementary Figure 8a**). SCA2 and SCA6 had a high number of differentially expressed RBPs within both the cerebellum and frontal cortex, potentially implying a specific role for RBPs in these two diseases. *HSPA6* is an RBP that is upregulated across several different ataxia subtypes. Interestingly, *ITPR1*, a gene with Purkinje cell type-specific expression^68^, has RBP function and is one of the most highly downregulated RBPs. These findings suggest the importance of RBP dysregulation in ataxia.

Given these findings, we identified differentially used introns. A total of 57,797 clusters were tested for each comparison. The total number of differentially spliced intron clusters and related genes (FDR p < 0.05) (**Figure 8a, b**) varied across disease, implying differential roles of splicing. This analysis also demonstrated that splicing dysregulation was largely tissue-specific with only 116 genes differentially spliced in both the frontal cortex and cerebellum. We explored whether differentially spiced intron clusters arose because of novel splicing events (those not in annotation). To achieve this, differentially spliced intron clusters (defined as those with FDR p < 0.05 and (|ΔPSI|) *≥* 0.1), were assigned to each category of splicing event (**Figure 8c**). We found that most of the differentially spliced intron clusters were annotated with reference to Gencode v.38 (60.3% to 70.1% across all comparison groups and regions). There were no significant differences between the proportions of the different types of junction annotation across different comparisons and the two brain regions (chi-square test FDR p > 0.05). Finally, very few genes were both differentially expressed and differentially spliced, independent of brain region or comparator groups (**Supplementary Figure 9**), indicating that the analysis of splicing was capturing an independent component of the pathophysiology. We annotated the junctions within the four causative genes (*FXN*, *ATXN1*, *ATXN2*, *CACNA1A*) with respect to the reference transcriptome. There was no statistically significant increase in the proportion of novel junction reads across each type of splicing event for any disease subtype within each of these Mendelian genes (**Figure 8d**). The only statistically significant finding was a higher proportion of novel donor junctions in *ATXN2* within the cerebellum of control individuals compared to disease (Kruskal Wallis FDR p = 0.0033) (**Figure 8d**). In the case of *FXN*, this likely reflects the downregulation and low overall coverage (mean (SD) read count across all samples: 267 (151)).

**Figure 8.**
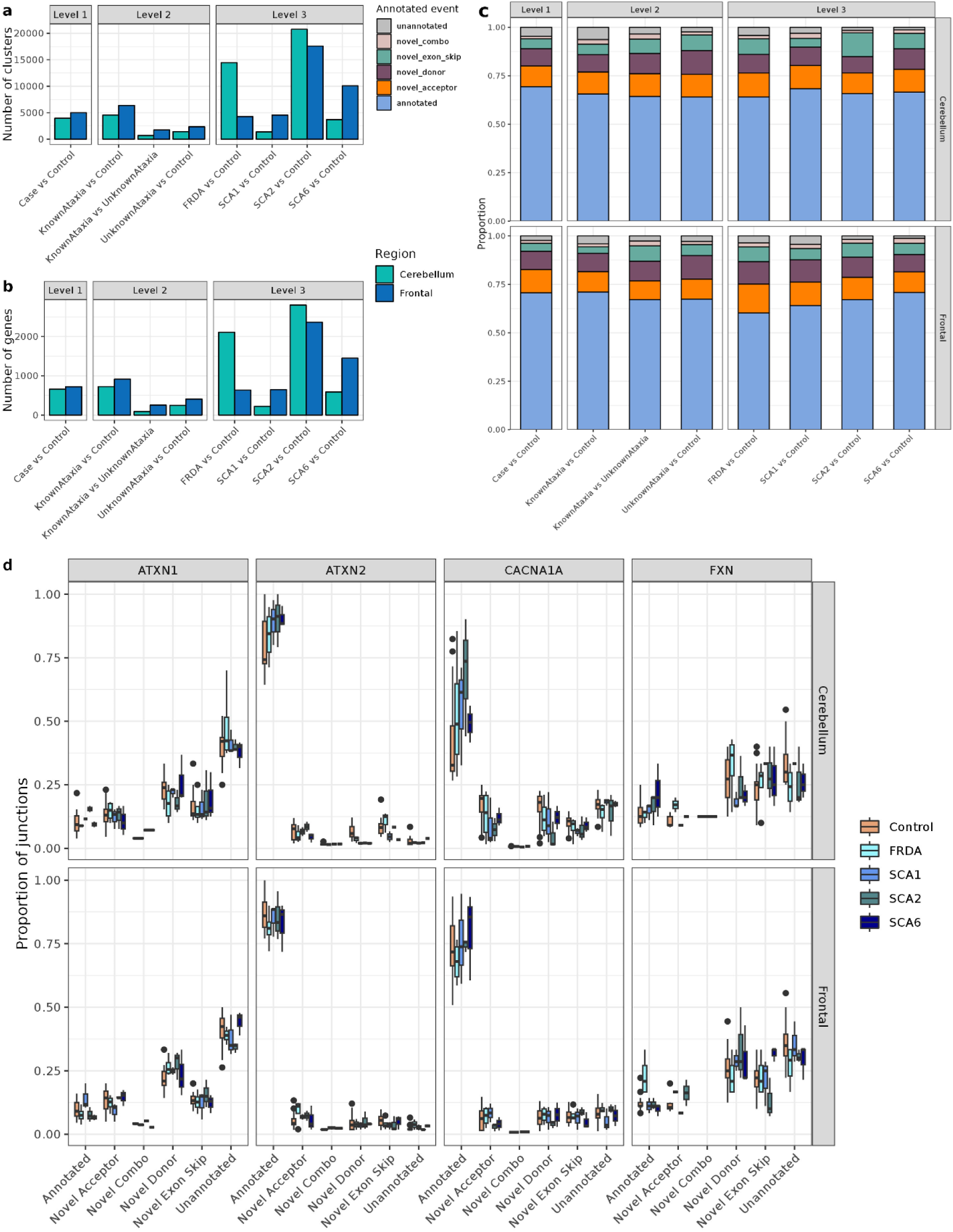
Annotation of junction reads. The number of significant (FDR p < 0.05 and (|ΔPSI|) *≥* 0.1) differentially spliced intron clusters (**a**) and genes into which the intron clusters fall (**b**) across the different comparator groups and brain regions. (**c**). The proportion of differentially spliced introns attributed to a particular splicing event category partitioned across different comparisons. “Novel combo” refers to novel combination events. (**d**). Comparison of the annotated splicing events for the junction reads (from STAR) in each disease group within the four causative Mendelian genes associated with SCA1, SCA2, SCA6 and FRDA. Unannotated group refers to no annotation at either the donor or acceptor sites.

### Pathway enrichment of differentially spliced genes

To identify the biological pathways which might be affected by splicing dysregulation in ataxia, each differentially spliced intron cluster was annotated to a gene using the reference transcriptome and gene set enrichment analysis was then performed. Pathway enrichment of differentially spliced genes showed that first, there was commonality in the affected pathways at a high annotation level within the cerebellum and frontal cortex (**Figure 9a**). Differentially spliced genes in both the cerebellum and frontal cortex were enriched for vesicle-transport, synapse; RNA-binding, nervous system development and actin cytoskeleton organisation (**Figure 9d**). The enrichment result for RNA-binding pathways was likely to be driven by the splicing of RBP genes themselves. In fact, up to 45% of differentially spliced genes in some groups (FRDA, SCA1, SCA2 cerebellum compared with controls) were genes associated with RBP function. Secondly, pathways related to nervous system development, focal adhesion were common across the comparator groups in both diagnosed and undiagnosed cases (**Figure 9b, e**); Lastly, whereas nervous system development and synaptic pathway enrichments were seen in SCA2 and SCA6 differentially spliced genes in frontal cortex, no enrichments of this kind were observed in FRDA and SCA2 individuals in either brain region. Instead, differentially spliced genes in the cerebellum of these cases were enriched for RNA-processing, protein transport, protein localisation to membrane, negative regulation of gene expression, cell cycle, actin cytoskeleton organisation and ubiquitin-dependent protein catabolic processes (**Figure 10c, f**). Interestingly, there was no evidence of cell type-specificity amongst differentially spliced genes. Thus, splicing dysregulation in ataxia is disease-specific.

**Figure 9.**
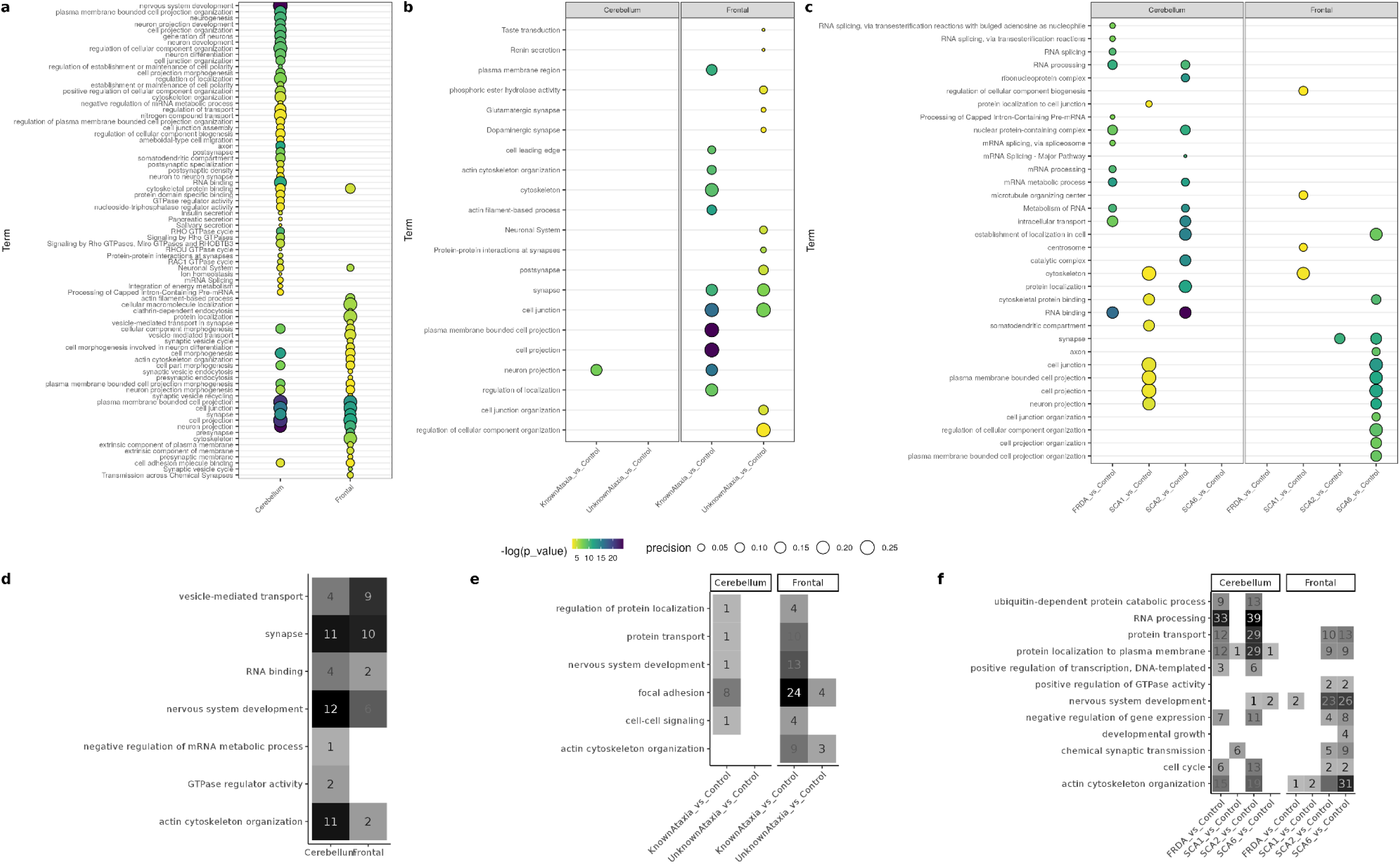
Pathway enrichment of differentially spliced genes. Pathway enrichment analysis and significantly enriched gene ontology (GO) terms associated with differentially spliced genes integrating all cases and controls (Level 1) (**a**); comparing cases with known and unknown molecular diagnosis (Level 2) (**b**); and comparing genetic subgroups (Level 3) (**c**). The most enriched pathways (defined by gSCS-corrected p value magnitude) are shown. Precision defines the proportion of genes in the input gene list that can be assigned a term. Corresponding tileplots (**d, e, f**) for Levels 1, 2, 3 analyses respectively showing semantically-summarised terms of all enriched GO terms. The numbers represent the number of terms summarised for each parent term

**Figure 10.**
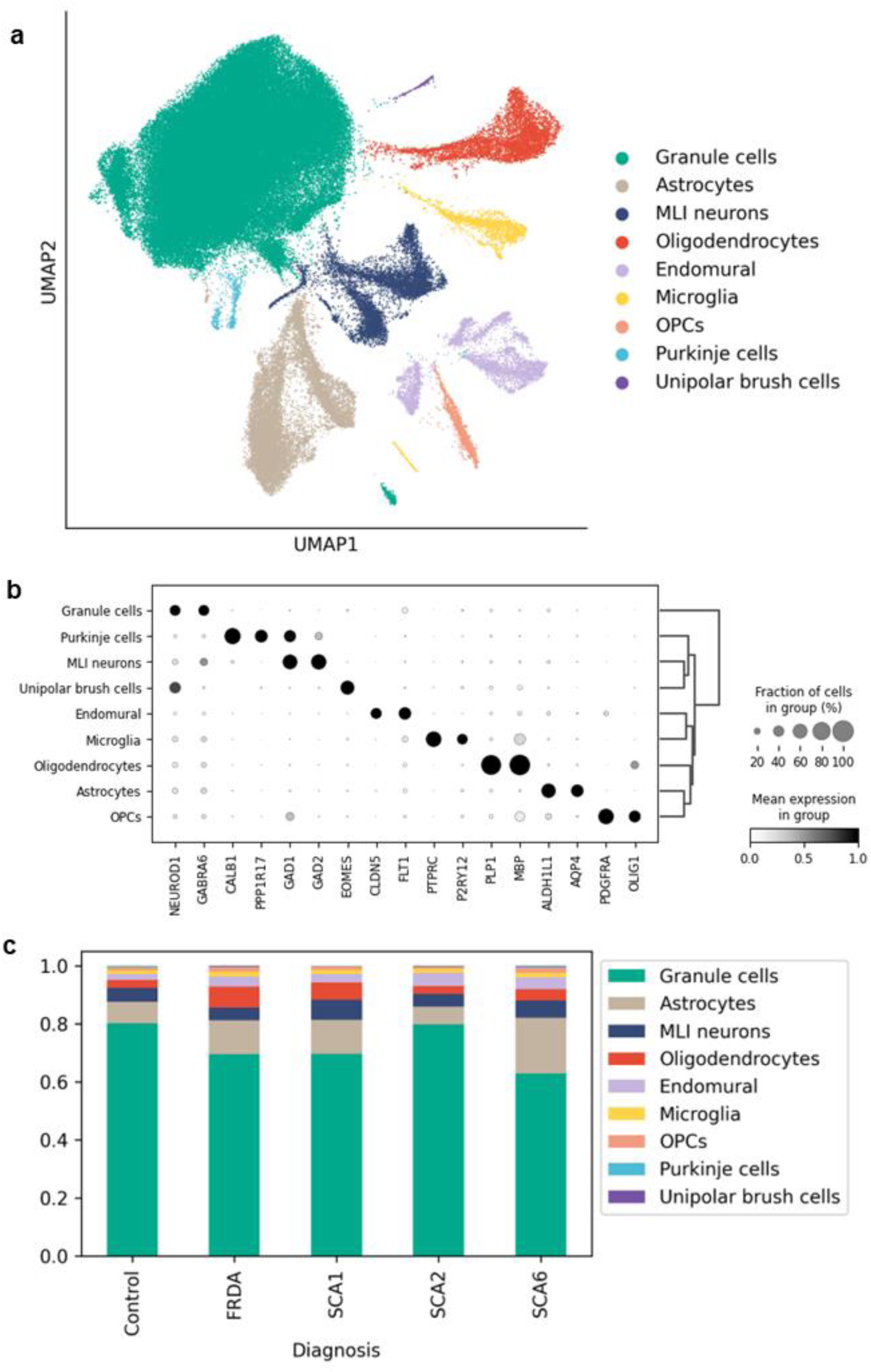
snRNA-sequencing data from cerebellar cortex. a) UMAP showing clustering of nuclei, labelled according to annotated cell types; b) Expression of cell type marker genes across nuclei clusters; c) Proportion of cell types across samples grouped by disease diagnosis.

### Single nuclear RNA-sequencing data supports a significant role for the innate immune system in disease

Cerebellar cortex samples from donors with SCA1, SCA2 and SCA6 together with matched controls underwent snRNA-sequencing. Nuclei were annotated to 9 major cell types: astrocytes, oligodendrocytes, oligodendrocyte precursor cells (OPCs), microglia, medium layer inter neurons (MLI neurons), Purkinje cells, unipolar brush cells, granule cells and endomural cells (**Figure 10a, b**). Although granule cells were the most abundant cell type across all diagnostic groups (**Figure 10c**), we were able to annotate Purkinje cells despite their relative loss in the disease state. Consistent with the neurodegenerative nature of the SCAs, we identified changes in cell type proportion in disease states. As expected across FRDA, SCA1 and SCA6, there was a lower proportion of neuronal cell types, including Purkinje cells, compared to the controls. However, disease-specific changes were also observed (**Figure 10c**). We noted that in SCA6, there was a lower proportion of granule cells, but a higher proportion of astrocytes. In contrast, in FRDA, there was a higher proportion of oligodendrocytes compared to matched controls.

Differential gene expression analysis was performed comparing each diagnosis group to control, as well as grouping all four ataxias together (denoted Ataxia vs. Control) (**Figure 11**). The highest number of differentially expressed genes (DEGs) was detected comparing all ataxia cerebellar samples to control in granule cells, likely reflecting the highest cell type proportion. Of note, SCA1 oligodendrocytes also displayed large amounts of differential expression compared to control samples, consistent with findings from the bulk RNA-sequencing data (**Figure 11a**). Across all cell types, the lowest numbers of DEGs was present when comparing FRDA to controls, again consistent with bulk RNA-sequencing results and supporting a loss-of-function without transcriptome-wide changes.

**Figure 11.**
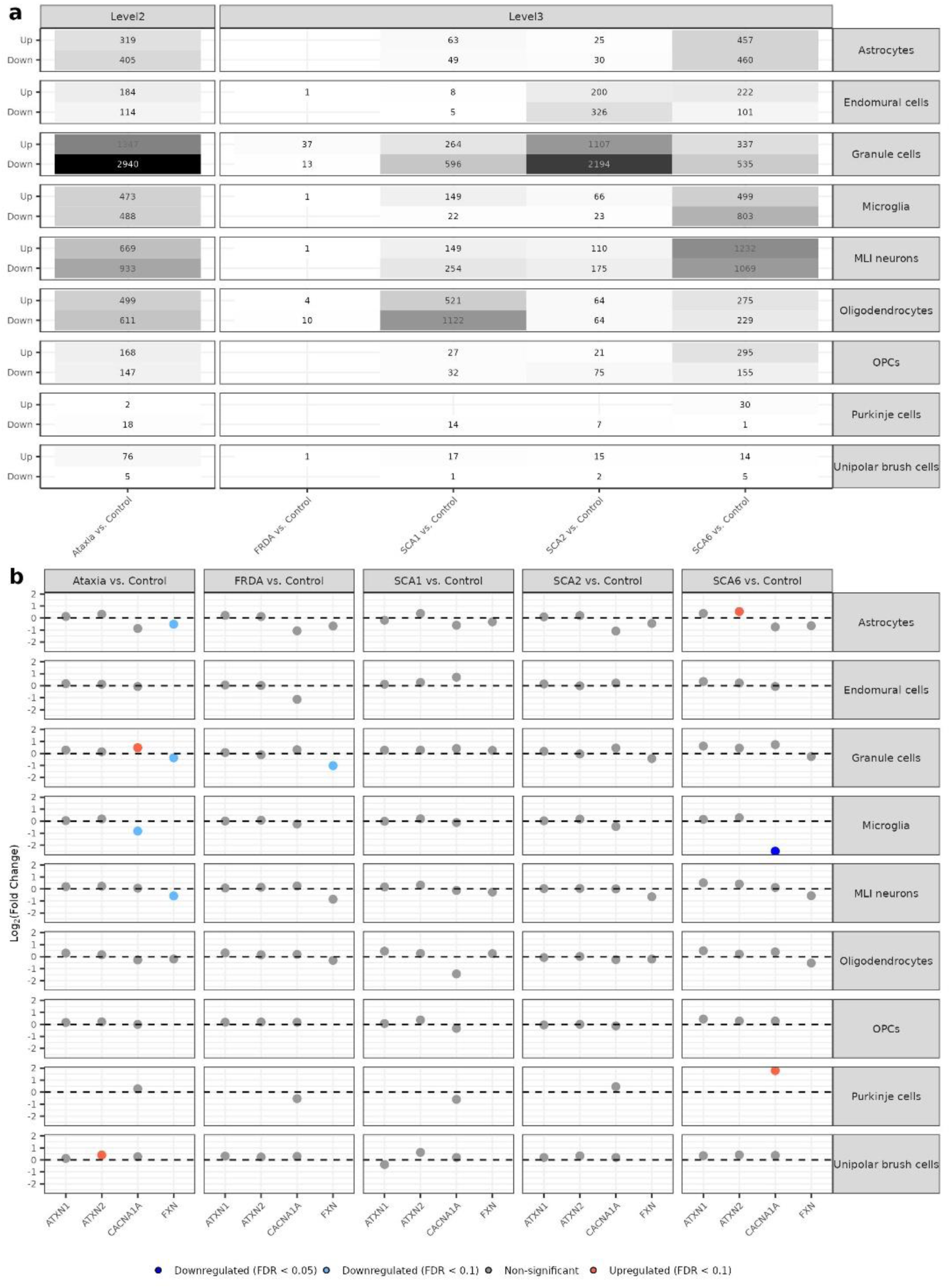
Differential gene expression analysis of snRNAseq data from cerebellar cortex. a) Total number of differentially expressed genes across each cell type and diagnosis. b) Change in expression of causative ataxia genes in each diagnosis comparison, across each cell type. “Ataxia” represents all cases combined.

When examining the expression changes of the Mendelian genes, *FXN* was downregulated in the granule cells of FRDA cerebellum. Interestingly, *CACNA1A* (the causative Mendelian gene for SCA6) was significantly upregulated in SCA6 Purkinje cells and downregulated in microglia (**Figure 11b**), a finding which required the acquisition of snRNAseq data.

Up- and downregulated DEGs were subsequently analysed for functional enrichment of GO terms (**Figure 12**). The highest number of enriched terms was detected in SCA6 compared to control microglia (**Figure 12a**). Upregulated terms in microglia included innate immune response (GO:0045087, FDR range = 4.34 × 10^−5^ – 3.65 × 10^−2^) and positive regulation of B-cell activation (GO:0050871, FDR range = 8.22 × 10^−12^ – 4.86 × 10^−2^) (**Figure 12b**). Commonalities in enriched pathways extended beyond microglia. In fact, we noted downregulated terms pertaining to transporter function and ubiquitous processes such as autophagy within neuronal cell types, again consistent with findings from bulk RNA-sequencing data.

**Figure 12.**
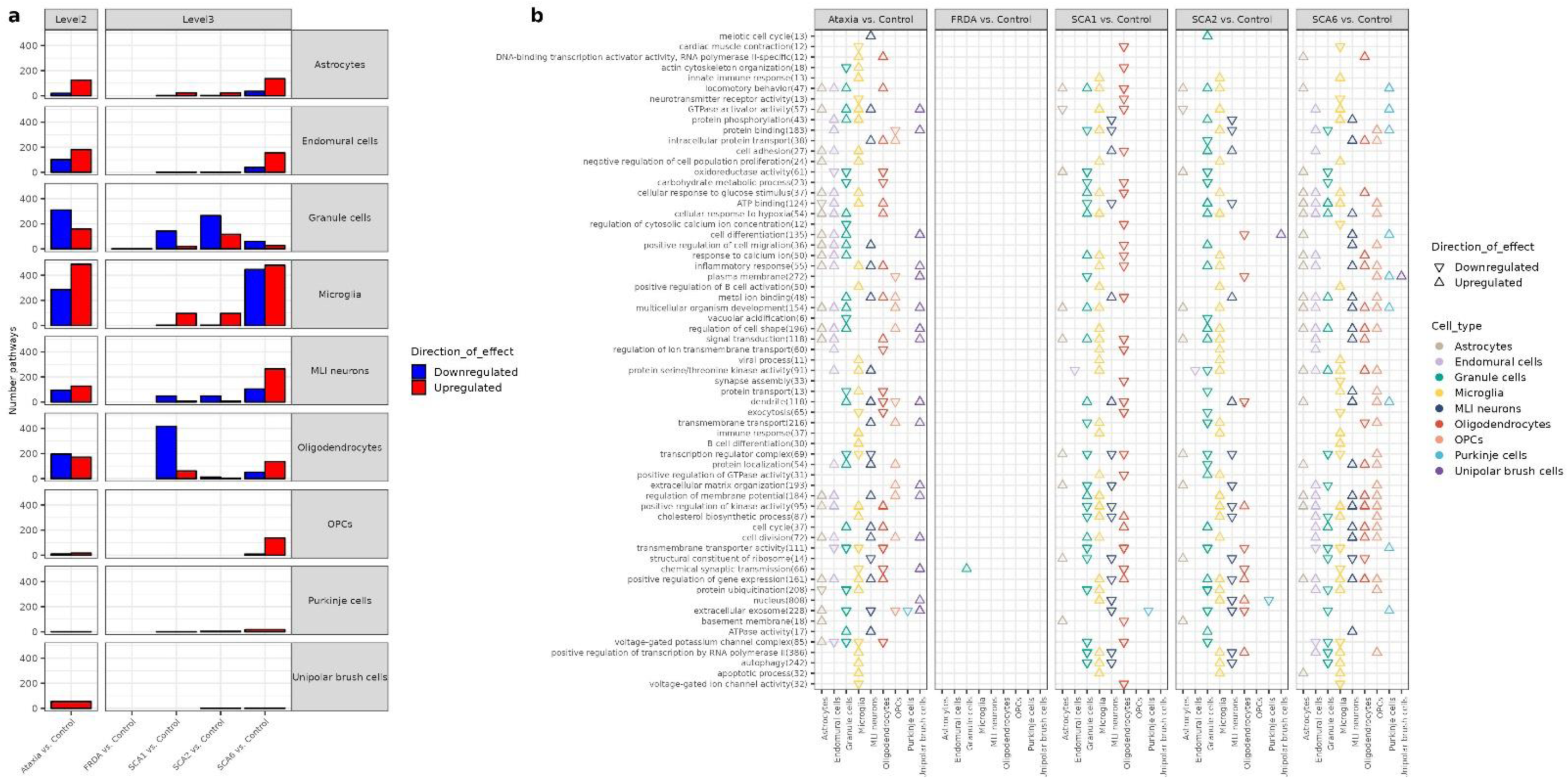
Functional enrichment analysis of genes differentially expressed across cerebellar cortex cell types. a) Total number of gene ontology terms enriched among differentially expressed genes (DEGs) in each diagnosis comparison and cell type. b) Gene ontology terms (reduced into parent terms, with the number of child terms indicated in the brackets) enriched across DEGs in each diagnosis comparison and cell type, excluding parent terms that are enriched in DEGs effected in both directions. The “Ataxia” comparison groups together FRDA, SCA1, SCA2 and SCA6 samples.

## Discussion

This study applied bulk and single nuclear RNA-sequencing to post-mortem brain tissue of donors with ataxia to gain further understanding of the underlying biology of disease. This analysis provided mechanistic insights that revealed that: (i) despite the commonalities in the genetics of ataxia, there were components of their transcriptional signatures which were distinct; (ii) there were extensive transcriptional changes evident not only in the cerebellum but also the frontal cortex in ataxia cases; (iii) activation of immune and inflammatory pathways, as well as involvement of non-neuronal cell types was a feature of all ataxias to a lesser or greater extent. Given the paucity of transcriptomic analyses of post-mortem human brain derived from individuals with ataxia, this study provides a novel resource to understand the mechanisms of disease in ataxia. Furthermore, taken together, these results highlight immune pathways as early and potentially important therapeutic targets, for further exploration.

All the molecularly diagnosed cases analysed in this study were repeat expansion disorders: SCAs are autosomal dominant exonic CAG repeat expansions and FRDA is an autosomal recessive GAA trinucleotide repeat expansion in intron-1 of the *FXN* gene. The existence of a common genetic lesion, particularly amongst the SCAs, made it tempting to imagine that these disorders might all be generated through a single and common pathogenic process arising from the repeat expansion itself. However, there were significant quantitative and qualitative differences in transcriptional responses across genotypes that were likely to be related to the genic context of the repeat expansion, the physiological gene function and the specific compensatory processes triggered. This was perhaps most evident when focusing on FRDA, where downregulation of *FXN* was the dominant change across both brain regions as previously reported^62,65,66^. Consistent with expectation, this was associated with downregulated heme processing pathways and upregulation of protein folding and apoptotic pathways in the cerebellum in keeping with the putative physiological function of frataxin^69^. Interestingly, while very few other differentially expressed genes were detected on a transcriptome-wide level, there was a relatively large number of differentially spliced intron clusters and RBPs in FRDA cerebellum. This is likely explained by the fact that GAA-repeats are known to bind to a multitude of different splicing factors^70^ and may also explain why at a transcriptome-wide level, there was evidence of an innate immune response in FRDA brain tissue with upregulation of *IL-1β* and *HSPA6* in the cerebellum and *IL-21R-AS1* in the frontal cortex.

Focusing on *ATXN2*, which is known to be a common mRNP granule component binding directly to RNA and other RBPs including TDP-43 and FUS^71^, widespread RBP dysregulation and differential splicing in both the cerebellum and frontal cortex was present, likely reflecting disruption of the physiological function of the gene. This was associated with a significant upregulation of genes involved in DNA methylation including the PRC2 pathway, again relating to the native role of ataxin-2 in miRNA-mediated gene silencing pathways or a compensatory mechanism to silence the repeat-containing DNA^72^. There was also a marked upregulation of genes involved in base excision repair, DNA damage and recruitment of ATM-mediated phosphorylation, pointing more clearly to compensatory processes that could explain commonalities across different forms of ataxia and the variable expressivity and progression of SCA2^73^. This was supported by differential expression of genes within the mismatch and base excision pathways across the brain in the disease state.

Consistent with this theme, there was a highly significant upregulation of *SPOCD1* in both frontal and cerebellar cortex of SCA6 patients compared to controls. SPOCD1 is a nuclear protein whose expression is normally restricted to the period of *de novo* genome methylation, and is essential for piRNA-directed methylation and silencing of young transposable elements including long interspersed nuclear element-1 (LINE1)^61^. This is of relevance as a recent study of ataxia telangiectasia found that LINE1 activation in cerebellar Purkinje cells was associated with progressive ataxia, coupled with a downregulation of transposable element regulators^25^. Dysregulation of transposable elements have been implicated in other neurological diseases^74^. Therefore, the upregulation in SCA6 of *SPOCD1*, a transposable element repressor, may imply a compensatory mechanism to LINE1 or transposable element activation in disease.

Importantly, transcriptional signatures of disease did not necessarily correlate in an obvious manner with the clinical syndromes of the affected patients. While cerebellar dysfunction is at the core of disease, ataxia syndromes differ in terms of their age-of-onset, clinical features, brain imaging and neuropathology. Pathology in FRDA is particularly widespread with atrophy of the dorsal root ganglia (DRG), sensory peripheral nerves, corticospinal tracts, and cerebellum^75^. While SCA 1, 2 and 17 are progressive ataxia with extra-cerebellar manifestations, SCA7 has additional retinal degeneration and SCA6 is a ‘pure’ ataxia syndrome with predominant cerebellar dysfunction^1^. Consequently, it would be expected that the greatest transcriptional dysregulation across all genotypes to be most apparent in the cerebellum. However, there was a comparable number of differentially expressed genes and pathways within both cerebellum and frontal cortex samples, especially within SCA1 and SCA6 affected individuals compared to controls. The extent of frontal involvement was particularly unexpected for SCA6, where the mild executive dysfunction, indicative of frontal cortical involvement would be subclinical or mild at best^76^. Focusing on SCA1 and SCA6, it was striking that the transcriptional changes in frontal cortex in cases were dominated by the upregulation of gene expression. Furthermore, these genes were highly enriched for inflammatory pathways as well as glial cell type-specific markers. Our snRNA-sequencing study showed a significant role for microglial activation in the SCA6 cerebellum and the role of oligodendrocytes in the SCA1 cerebellum. The latter finding is consistent with a previous study showing transcriptional dysregulation in mouse and human SCA1 cerebellar oligodendroglia^24^. These findings highlighted the extent of cortical involvement in ataxia at the transcriptional level, something which is not apparent from current clinical, pathological, and radiological data. This suggests that inflammatory biomarkers such as PET imaging and blood might be a means of tracking the disease progress.

These observations also pointed to the unexpected importance of inflammation and innate immune function across all ataxia cases and genetic subtypes (**Figure 13**). While there is growing awareness of inflammation as an important feature of other neurodegenerative disorders, with a wealth of evidence in Alzheimer’s and Parkinson’s diseases in particular (reviewed in ^77–80^), the roles of inflammation and non-neuronal cells are not well characterised or recognised in ataxia. In neurodegeneration, damage-associated molecular patterns (DAMPs) are endogenous signals released in the event of cellular stress to misfolded and aggregated proteins such as alpha-synuclein in Parkinson’s disease, and trigger the innate immune system by interacting with pattern recognition receptors (PRRs)^80^. Given that samples originate from individuals with longstanding disease and the cerebellum is affected much earlier than the frontal cortex across all forms of ataxia, the strong inflammatory signals in the frontal cortex may be representative of the initial disease phase. Interestingly, in SCA6, a late-onset slowly progressive ataxia, there were clear transcriptomic signatures relating to upregulation of the innate immune system in the frontal cortex but both innate and adaptive responses in the cerebellum, again suggesting progression. Furthermore, an early role for the innate immune system in ataxia is independently supported *in vivo* through the observation of increases in peripheral immune markers in SCA2 preclinical carriers^81^ and mouse models suggesting that attenuating immune responses could be beneficial. Previous studies of the knock-in SCA6 mouse suggested that microglial activation preceded the onset of Purkinje cell loss in the cerebellum and that genetically ablating MyD88 (part of the TLR signalling pathway and a key component of innate immune receptor signalling as TLR is a type of PRR) partially rescued Purkinje cell loss^82^. Similarly, a recent study using the SCA1 mouse model found that inhibiting JNK-dependent glial activation in the cerebellum reduces inflammation and improves the corresponding phenotype and pathology, again suggesting a causal role for inflammation in disease^83^.

**Figure 13.**
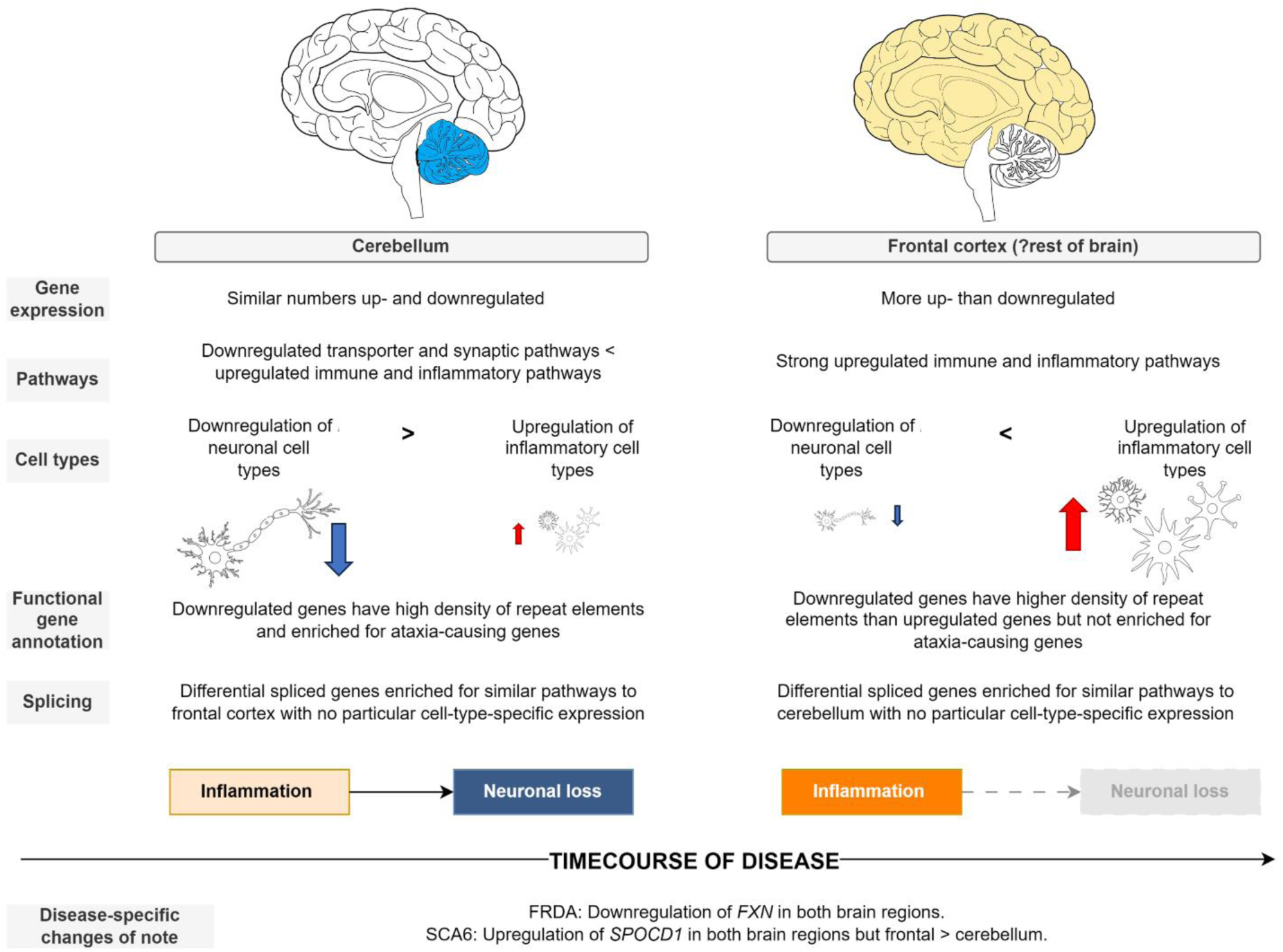
Schematic summarising findings from ataxia RNA-sequencing. There are differential signatures of gene expression and associated pathway enrichment, cell type-specific enrichment and functional gene annotation, as well as within differential splicing across the two brain regions. The overall signature is a pattern of more profound neuronal loss than inflammation within the cerebellum in ataxia, compared to a less severe signature of neuronal loss compared to stronger inflammatory signals in the frontal cortex.

Even if innate immune activation is an early event in ataxia, the triggers and amplifiers of that process remain unclear. Extrapolating from the example of Huntington’s disease, where the CAG repeat and expanded polyglutamine protein has been shown to directly trigger inflammation in the absence of endogenous signaling of cellular damage^84^, repeat expansions might be an obvious candidate in the context of SCAs. More broadly across neurodegenerative diseases, there is an increasing recognition of the potential for endogenous signals released in the event of stress to trigger the innate immune system by interacting with PRRs^80,85^. Misfolded or aggregated proteins, heat shock proteins (HSP), DNA fragments and ATP have all been reported to be DAMPs recognised by PRRs and inducing the expression of pro-inflammatory cytokines including IL-6, IL-8, IL-12, TNF*α*, IFN*β*, TGF*β*^86^. This analysis across ataxia genotypes is consistent with this process with, e.g., *CXCL8* (IL-8) upregulation in SCA6. IL-6 and IL-8 have specific pro-inflammatory involvement via the DAMPs signaling to trigger microglial activation^86^ and were found to be upregulated in the ataxia cerebellum. Additionally, strongly upregulated expression of *HSP* including *HSPA6* in ataxia cases compared to controls could reflect DAMPs from neuronal damage, which have an important role in triggering innate immune reaction in neurodegeneration^86^.

Another potential inflammatory trigger is the expression of abnormal RNA species and even proteins generated through splicing dysregulation^87^. Few studies on ataxia have investigated the transcriptome-wide level changes in splicing despite several of the SCA genes being implicated in transcription, and the potential for aberrant splicing due to transcriptional stalling from the repeat expansion and/or sequestration of essential splicing factors in RNA foci^88^. Investigations of splicing events and differential splicing within the causative Mendelian genes revealed the high complexity of splicing events with difficulties inferring pathogenic mechanisms using short-read RNA-sequencing data alone. Beyond the causative genes, differential splicing was prevalent in ataxia on a transcriptome-wide level and genes with differentially spliced intron clusters were different to those that were differentially expressed. While there were clear regional differences in cell type-specific expression and functional pathway enrichment in differentially expressed genes, these features were not appreciable in those differentially spliced genes, which were largely enriched for RNA-processing and protein transport pathways. Consequently, splicing dysregulation may be insufficient to explain vulnerability of the cerebellum specifically in ataxias.

There are several limitations of this study that provide motivation for future work. First, given that ataxias are rare disorders with clinical and genetic heterogeneity, poor diagnostic yield, and a resulting paucity of post-mortem specimens, this cohort used was relatively small compared to larger RNA-sequencing studies of common neurodegenerative conditions. However, the number of differentially expressed genes increased with smaller sample sizes that focused on specific genotypes highlighting the heterogeneity of data. The number of samples of ataxia of each genetic subtype was comparable to the numbers used for analysis in other RNA-sequencing studies of ataxia^25^. Leafcutter does not allow modelling of intron retention events which could potentially be important in the pathogenic mechanism of repeat expansion disorders^89^. Understanding the full splicing map in disease is crucial given that splicing patterns can be used to personalise splice-switching therapy in ataxia^90^ and that previous studies have shown annotation of transcripts is far from complete^91^. Thus, using long-read RNA sequencing would be helpful in further understanding transcript expression.

## Conclusion

This transcriptomic study revealed unexpected immune and inflammatory upregulation in ataxia, across the molecularly diagnosed subtypes and undiagnosed cases. This provides key insights into the biology of disease in repeat expansion disorders, and highlights the complexities and importance of non-neuronal cells in disease. It further identified molecular signatures of disease between different genetic subtypes and across the brain regions, and is the one of the first to simultaneously assess both the transcriptome-wide impact of splicing and gene expression in ataxia. In an era of much promise provided by RNA-based therapies for neurological disease, this study supports the need for detailed transcriptomic mapping of disease states to inform full understanding of the biology of disease at the RNA level.

## Acknowledgments

We are grateful to those individuals who have donated brain tissue and their relatives to the Victoria Brain Bank (Australia) and Queen Square Brain (UK), without whom this work would not have been possible. We are grateful to Queen Square Brain Bank and Victoria Brain Bank staff for their assistance with donation and material preparation process. We are grateful to the following funders who supported this work: Wellcome Trust, Medical Research Council (MRC) UK, MSA Trust, National Institute for Health Research University College London Hospitals Biomedical Research Centre (NIHR-BRC), Michael J Fox Foundation (MJFF), Fidelity Trust, Dolby Family Fund, Rosetrees Trust, British Embassy Gulf Strategy Fund, Alzheimer’s Research UK (ARUK), MSA Coalition, NIH NeuroBioBank and MRC Brain Bank Network.

