## Supplementary for "Transcriptomic analysis of repeat expansion-ataxias uncovers distinct non-neuronal cell type-specific signatures of disease across the human brain"

Chen *et al.* 2025

### Supplementary Materials

i. Supplementary Tables
ii. Supplementary Figures
iii. Supplementary Methods

### Supplementary Tables

| Syndrome | Clinical Features | Age-of-onset | Gene | Mode of inheritance | Repeat motif | Putative gene function | Gain-of-function | Loss-of-function | RNA-binding protein seques-tration | RNA foci | RAN transla-tion |
|---|---|---|---|---|---|---|---|---|---|---|---|
| **SCA1** | Ataxia with pyramidal involvement and cognitive impairment | Bimodal ~30 to 40 years of age, but a proportion have a more pure ataxia with onset >60 years | *ATXN1* | AD | CAG | Transcription regulation | + | + | ? | + | ? |
| **SCA2** | Ataxia with ophthalmoparesis, parkinsonism, pyramidal features | Fourth decade | *ATXN2* | AD | CAG | RNA processing | + | + | + | + | + |
| **SCA6** | Slowly progressive, 'pure' ataxia | After fourth decade | *CACNA1A* | AD | CAG | Voltage-gated calcium channel | + | ? | ? | + | ? |
| **SCA7** | Ataxia with cone-rod retinal dystrophy | Range from adolescence to adult onset. In infantile onset cases, disease is severe with multiorgan failure | *ATXN7* | AD | CAG | Part of transcription co-activator complex, nuclear-cytoplasmic shuttling | + | ? | ? | + | ? |
| **SCA17** | Ataxia, dementia, involuntary movements such as chorea and dystonia | Usually by age 50 years | *TBP* | AD | CAG | Transcription initiation factor | + | + | ? | + | ? |
| **FRDA** | Slowly progressive ataxia with pyramidal features, scoliosis, bladder dysfunction and loss of position and vibration sensation | Usually by 25 years but a quarter have adult-onset disease | *FXN* | AR | GAA | Regulating mitochondrial iron transport | ? | + | ? | - | ? |

**Supplementary Table 1. Summary of mechanisms of disease in ataxia associated with repeat expansion disorders.** Summary of different categories of disease mechanisms in relation to common spinocerebellar ataxias (SCA) and Friedreich's ataxia (FRDA) associated with repeat expansion disorders in this study. Clinical feature an age-of-onset information are taken from [64]. For mode of inheritance, AD represents autosomal dominant and AR represents autosomal recessive. + indicates presence of a mechanism. - indicates absence of a mechanism. ? indicates unknown presence of mechanism.

**Supplementary Table 2. Demographics from brain donors.** Anonymised IDs for samples are provided. Samples are partitioned according to Level 1, Level 2 and Level 3 categories. Diagnosed ataxia refers to molecularly diagnosed cases and undiagnosed ataxia refers to cases without current molecular diagnosis. PMI refers to post-mortem interval. RIN refers to RNA integrity number. For brain banks, QSBB = Queen Square Brain Bank and VBB = the Victorian Brain Bank.

| ID | Level 1 | Level 2 | Level 3 | PMI (hours) | RIN | Brain region | Brain Bank | Age at Death (years) | Sex |
|---|---|---|---|---|---|---|---|---|---|
| CO10A | Control | Control | Control | 79 | 3.7 | Cerebellum | QSBB | 76 | M |
| CO10A | Control | Control | Control | 79 | 6.5 | Frontal | QSBB | 76 | M |
| CO11A | Control | Control | Control | 60 | 4.4 | Cerebellum | QSBB | 96 | F |
| CO11A | Control | Control | Control | 60 | 5.2 | Frontal | QSBB | 96 | F |
| CO12A | Control | Control | Control | 76 | 1.5 | Cerebellum | QSBB | 71 | F |
| CO12A | Control | Control | Control | 76 | 3.2 | Frontal | QSBB | 71 | F |
| CO13A | Control | Control | Control | 54 | 6.8 | Cerebellum | QSBB | 84 | F |
| CO13A | Control | Control | Control | 54 | 6 | Frontal | QSBB | 84 | F |
| CO14B | Control | Control | Control | 50.5 | 7.7 | Cerebellum | VBB | 38.2 | F |
| CO14B | Control | Control | Control | 50.5 | 7.5 | Frontal | VBB | 38.2 | F |
| CO15B | Control | Control | Control | 56 | 7.6 | Cerebellum | VBB | 32.3 | F |
| CO15B | Control | Control | Control | 56 | 7.7 | Frontal | VBB | 32.3 | F |
| CO16B | Control | Control | Control | 52 | 6.9 | Cerebellum | VBB | 38.8 | F |
| CO16B | Control | Control | Control | 52 | 7.6 | Frontal | VBB | 38.8 | F |
| CO17B | Control | Control | Control | 33 | 6.5 | Cerebellum | VBB | 52.1 | M |
| CO17B | Control | Control | Control | 33 | 6.9 | Frontal | VBB | 52.1 | M |
| CO18B | Control | Control | Control | 43 | 7.5 | Cerebellum | VBB | 84.9 | M |
| CO18B | Control | Control | Control | 43 | 6.1 | Frontal | VBB | 84.9 | M |
| CO19B | Control | Control | Control | 50 | 3.1 | Cerebellum | VBB | 49 | M |

| ID | Level 1 | Level 2 | Level 3 | PMI (hours) | RIN | Brain region | Brain Bank | Age at Death (years) | Sex |
|---|---|---|---|---|---|---|---|---|---|
| CO19B | Control | Control | Control | 50 | 6.7 | Frontal | VBB | 49 | M |
| CO1A | Control | Control | Control | 14 | 6 | Cerebellum | QSBB | 81 | F |
| CO1A | Control | Control | Control | 14 | 5.7 | Frontal | QSBB | 81 | F |
| CO20B | Control | Control | Control | 64 | 6.8 | Cerebellum | VBB | 52 | M |
| CO20B | Control | Control | Control | 64 | 6.5 | Frontal | VBB | 52 | M |
| CO21B | Control | Control | Control | 46 | 7.1 | Cerebellum | VBB | 76 | M |
| CO22B | Control | Control | Control | 36.5 | 5.8 | Cerebellum | VBB | 81 | M |
| CO22B | Control | Control | Control | 36.5 | 5.3 | Frontal | VBB | 81 | M |
| CO23B | Control | Control | Control | 71 | 7.1 | Cerebellum | VBB | 69.6 | M |
| CO23B | Control | Control | Control | 71 | 6.6 | Frontal | VBB | 69.6 | M |
| CO2A | Control | Control | Control | NA | 6.3 | Cerebellum | QSBB | 79 | M |
| CO2A | Control | Control | Control | NA | 6.1 | Frontal | QSBB | 79 | M |
| CO3A | Control | Control | Control | 40 | 6.8 | Cerebellum | QSBB | 81 | M |
| CO3A | Control | Control | Control | 40 | 6.7 | Frontal | QSBB | 81 | M |
| CO4A | Control | Control | Control | 59 | 6.7 | Cerebellum | QSBB | 80 | F |
| CO4A | Control | Control | Control | 59 | 5.9 | Frontal | QSBB | 80 | F |
| CO5A | Control | Control | Control | 51 | 4.4 | Cerebellum | QSBB | 81 | M |
| CO5A | Control | Control | Control | 51 | 6.5 | Frontal | QSBB | 81 | M |
| CO6A | Control | Control | Control | 79 | 5.5 | Cerebellum | QSBB | 79 | F |

| ID | Level 1 | Level 2 | Level 3 | PMI (hours) | RIN | Brain region | Brain Bank | Age at Death (years) | Sex |
|---|---|---|---|---|---|---|---|---|---|
| CO6A | Control | Control | Control | 79 | 7.1 | Frontal | QSBB | 79 | F |
| CO7A | Control | Control | Control | 49 | 2.5 | Cerebellum | QSBB | 80 | F |
| CO7A | Control | Control | Control | 49 | 5.7 | Frontal | QSBB | 80 | F |
| CO8A | Control | Control | Control | 105 | 3.7 | Cerebellum | QSBB | 79 | M |
| CO8A | Control | Control | Control | 105 | 5.7 | Frontal | QSBB | 79 | M |
| CO9A | Control | Control | Control | 46.5 | 6.6 | Cerebellum | QSBB | 90 | M |
| CO9A | Control | Control | Control | 46.5 | 6.3 | Frontal | QSBB | 90 | M |
| C10A | Case | Diagnosed Ataxia | SCA6 | 45 | 4.2 | Cerebellum | QSBB | 95 | F |
| C10A | Case | Diagnosed Ataxia | SCA6 | 45 | 5.2 | Frontal | QSBB | 95 | F |
| C11A | Case | Diagnosed Ataxia | SCA17 | 52.3 | 3.4 | Cerebellum | QSBB | 66 | M |
| C11A | Case | Diagnosed Ataxia | SCA17 | 52.3 | 6.6 | Frontal | QSBB | 66 | M |
| C12A | Case | Diagnosed Ataxia | SCA1 | 55 | 5.9 | Cerebellum | QSBB | 61 | M |
| C12A | Case | Diagnosed Ataxia | SCA1 | 55 | 7.5 | Frontal | QSBB | 61 | M |
| C13A | Case | Diagnosed Ataxia | SCA1 | 40 | 2.8 | Cerebellum | QSBB | 71 | F |
| C13A | Case | Diagnosed Ataxia | SCA1 | 40 | 5.2 | Frontal | QSBB | 71 | F |
| C14A | Case | Diagnosed Ataxia | FRDA | 66 | 4.7 | Cerebellum | QSBB | 76 | F |
| C14A | Case | Diagnosed Ataxia | FRDA | 66 | 6.5 | Frontal | QSBB | 76 | F |
| C15A | Case | Diagnosed Ataxia | FRDA | 48 | 2.8 | Cerebellum | QSBB | 37 | M |
| C15A | Case | Diagnosed Ataxia | FRDA | 48 | 5.5 | Frontal | QSBB | 37 | M |

| ID | Level 1 | Level 2 | Level 3 | PMI (hours) | RIN | Brain region | Brain Bank | Age at Death (years) | Sex |
|---|---|---|---|---|---|---|---|---|---|
| C17B | Case | Diagnosed Ataxia | SCA6 | 60 | 3.6 | Cerebellum | VBB | 81.3 | M |
| C17B | Case | Diagnosed Ataxia | SCA6 | 60 | 5.9 | Frontal | VBB | 81.3 | M |
| C18B | Case | Diagnosed Ataxia | SCA6 | 19.5 | 5.8 | Cerebellum | VBB | 85.5 | M |
| C18B | Case | Diagnosed Ataxia | SCA6 | 19.5 | 7.1 | Frontal | VBB | 85.5 | M |
| C19B | Case | Diagnosed Ataxia | SCA2 | 38 | 2.9 | Cerebellum | VBB | 38.2 | F |
| C19B | Case | Diagnosed Ataxia | SCA2 | 38 | 6.2 | Frontal | VBB | 38.2 | F |
| C20B | Case | Diagnosed Ataxia | SCA2 | 5.5 | 5.4 | Cerebellum | VBB | 38.9 | F |
| C20B | Case | Diagnosed Ataxia | SCA2 | 5.5 | 7.8 | Frontal | VBB | 38.9 | F |
| C21B | Case | Diagnosed Ataxia | SCA2 | 77.5 | 2.8 | Cerebellum | VBB | 51.9 | M |
| C21B | Case | Diagnosed Ataxia | SCA2 | 77.5 | 6.3 | Frontal | VBB | 51.9 | M |
| C22B | Case | Diagnosed Ataxia | SCA17 | 17 | 4.1 | Cerebellum | VBB | 52.6 | F |
| C22B | Case | Diagnosed Ataxia | SCA17 | 17 | 6.4 | Frontal | VBB | 52.6 | F |
| C23B | Case | Diagnosed Ataxia | SCA1 | 12 | 4.8 | Cerebellum | VBB | 69.7 | M |
| C23B | Case | Diagnosed Ataxia | SCA1 | 12 | 7.4 | Frontal | VBB | 69.7 | M |
| C24B | Case | Diagnosed Ataxia | FRDA | 45 | 5.4 | Cerebellum | VBB | 32.9 | F |
| C24B | Case | Diagnosed Ataxia | FRDA | 45 | 7.4 | Frontal | VBB | 32.9 | F |
| C25B | Case | Diagnosed Ataxia | FRDA | 65.5 | 7.3 | Cerebellum | VBB | 50.8 | M |
| C25B | Case | Diagnosed Ataxia | FRDA | 65.5 | 8.1 | Frontal | VBB | 50.8 | M |
| C9A | Case | Diagnosed Ataxia | SCA7 | 27 | 6.6 | Cerebellum | QSBB | 88 | F |

| ID | Level 1 | Level 2 | Level 3 | PMI (hours) | RIN | Brain region | Brain Bank | Age at Death (years) | Sex |
|---|---|---|---|---|---|---|---|---|---|
| C9A | Case | Diagnosed Ataxia | SCA7 | 27 | 4.8 | Frontal | QSBB | 88 | F |
| C2A | Case | Undiagnosed Ataxia | Undiagnosed Ataxia | 38 | 4.9 | Cerebellum | QSBB | 66 | F |
| C2A | Case | Undiagnosed Ataxia | Undiagnosed Ataxia | 38 | 5.4 | Frontal | QSBB | 66 | F |
| C3A | Case | Undiagnosed Ataxia | Undiagnosed Ataxia | 27.5 | 4 | Cerebellum | QSBB | 56 | M |
| C3A | Case | Undiagnosed Ataxia | Undiagnosed Ataxia | 27.5 | 7 | Frontal | QSBB | 56 | M |
| C4A | Case | Undiagnosed Ataxia | Undiagnosed Ataxia | 21 | 1.9 | Cerebellum | QSBB | 77 | M |
| C4A | Case | Undiagnosed Ataxia | Undiagnosed Ataxia | 21 | 2.2 | Frontal | QSBB | 77 | M |
| C5A | Case | Undiagnosed Ataxia | Undiagnosed Ataxia | 9 | 5.2 | Cerebellum | QSBB | 70 | F |
| C5A | Case | Undiagnosed Ataxia | Undiagnosed Ataxia | 9 | 7.8 | Frontal | QSBB | 70 | F |
| C6A | Case | Undiagnosed Ataxia | Undiagnosed Ataxia | 66 | 3.3 | Cerebellum | QSBB | 66 | F |

| ID | Level 1 | Level 2 | Level 3 | PMI (hours) | RIN | Brain region | Brain Bank | Age at Death (years) | Sex |
|---|---|---|---|---|---|---|---|---|---|
| C6A | Case | Undiagnosed Ataxia | Undiagnosed Ataxia | 66 | 6.8 | Frontal | QSBB | 66 | F |
| C7A | Case | Undiagnosed Ataxia | Undiagnosed Ataxia | 26 | 3.6 | Cerebellum | QSBB | 75 | M |
| C7A | Case | Undiagnosed Ataxia | Undiagnosed Ataxia | 26 | 4.4 | Frontal | QSBB | 75 | M |
| C8A | Case | Undiagnosed Ataxia | Undiagnosed Ataxia | 103 | 7.5 | Cerebellum | QSBB | 56 | M |
| C8A | Case | Undiagnosed Ataxia | Undiagnosed Ataxia | 103 | 7.1 | Frontal | QSBB | 56 | M |

**Supplementary Table 3. Cell type descriptions of Allen Brain Atlas data.** Description of cell types used for expression-weighted cell type enrichment analysis using Allen Brain Atlas data.

| *Cell type* | *Description* | *Major Class* |
|---|---|---|
| **Astrocyte** | Astrocyte | Astrocyte |
| **GABAergic_LAMP5** | Lysosomal Associated Membrane Protein Family Member 5 (LAMP5)-expressing GABAergic neuron | Inhibitory neuron |
| **GABAergic_PAX6** | Paired box 6 (PAX6)-expressing GABAergic neuron | Inhibitory neuron |
| **GABAergic_PVALB** | Parvalbumin (PVALB)-expressing GABAergic neuron | Inhibitory neuron |
| **GABAergic_SST** | Somatostatin (SST)-expressing GABAergic neuron | Inhibitory neuron |
| **GABAergic_VIP** | Vasoactive intestinal peptide (VIP)-expressing GABAergic neuron | Inhibitory neuron |
| **Glutamatergic_IT** | Glutamatergic intratelencephalic neurons | Excitatory neuron |
| **Glutamatergic_L4_IT** | Glutamatergic intratelencephalic neurons Layer 4 | Excitatory neuron |
| **Glutamatergic_L5_6_IT_Car3** | Glutamatergic intratelencephalic neurons Layer 5/6 | Excitatory neuron |
| **Glutamatergic_L5_6_NP** | Glutamatergic near-projecting neurons Layer 5/6 | Excitatory neuron |
| **Glutamatergic_L5_ET** | Glutamatergic extratelencephalic projecting neurons Layer 5 | Excitatory neuron |
| **Glutamatergic_L6_CT** | Glutamatergic corticothalamic neurons Layer 6 | Excitatory neuron |
| **Glutamatergic_L6b** | Glutamatergic neurons Layer 6b | Excitatory neuron |
| **Microglia** | Microglia | Microglia |
| **Oligodendrocyte** | Oligodendrocyte | Oligodendrocyte |
| **OPC** | Oligodendrocyte precursor cell | Oligodendrocyte |
| **Vascular cells** | Vascular cells | Non-neuronal |

**Supplementary Table 4. Gene markers used for cluster cell type annotation.**

| Major cell type | Gene markers | References |
|---|---|---|
| Astrocytes | *ALDH1L1,*<br>*AQP4* | https://www.cell.com/neuron/fulltext/S0896-6273(23)00844-9 |
| Endomural cells | *CLDN5,*<br>*FLT1* | https://www.biorxiv.org/content/10.1101/2022.09.26.509462v1<br>https://www.cell.com/neuron/fulltext/S0896-6273(23)00844-9 |
| Granule cells | *NEUROD1,*<br>*GABRA6* | https://www.cell.com/neuron/fulltext/S0896-6273(23)00844-9 |
| Microglia | *PTPRC,*<br>*P2RY12* | https://www.nature.com/articles/ncomms11295<br>https://www.cell.com/neuron/fulltext/S0896-6273(23)00844-9 |
| MLI neurons | *GAD1,*<br>*GAD2* | https://www.cell.com/neuron/fulltext/S0896-6273(23)00844-9 |
| Oligodendrocytes | *PLP1,*<br>*MBP* | https://www.biorxiv.org/content/10.1101/2022.09.26.509462v1 |
| Oligodendrocyte precursor cells | *PDGFRA,*<br>*OLIG1* | https://www.cell.com/neuron/fulltext/S0896-6273(23)00844-9 |
| Purkinje cells | *CALB1,*<br>*PPP1R17* | https://www.cell.com/neuron/fulltext/S0896-6273(23)00844-9 |
| Unipolar brush cells | *EOMES* | https://www.cell.com/neuron/fulltext/S0896-6273(23)00844-9 |

**Supplementary Table 5. Nuclei count by cell type**

| Major cell type | Number of nuclei |
|---|---|
| Astrocytes | 12508 |
| Endomural cells | 3961 |
| Granule cells | 101037 |
| Microglia | 1898 |
| MLI neurons | 6674 |
| Oligodendrocytes | 5173 |
| Oligodendrocyte precursor cells | 1565 |
| Purkinje cells | 399 |
| Unipolar brush cells | 227 |

**Supplementary Table 6. Sample comparisons analysed in differential expression analysis of snRNA-sequencing data.** Sample groups are labelled by diagnosis group and brain region, separated by underscores.

| Contrast | Sample group comparison |
|---|---|
| SCA1 vs. control | group_SCA1 – group_Control |
| SCA2 vs. control | group_SCA2 – group_Control" |
| SCA6 vs. control | group_SCA6 – group_Control" |
| FRDA vs. control | group_FRDA – group_Control" |
| Ataxia vs. control | (group_SCA1 + group_SCA2 + group_SCA6 + group_FRDA)/4 – group_Control |

**Supplementary Table 7. Comparison of sample measures.** Sample demographics and measures are compared across the brain regions between cases and controls. Continuous variables are compared using the Wilcoxon rank sum test and categorical variables using the chi-squared test. SD refers to the standard deviation of the mean and n refers to the total number of samples within that category. P value of the statistical test with significant differences of $p < 0.05$ denoted by the asterisk. PMI refers to the post-mortem interval and RIN refers to the RNA integrity number of the sample.

| Sample measure | Region | n (case) | n (control) | Case: Mean (SD) | Control: Mean (SD) | P value | Test |
|---|---|---|---|---|---|---|---|
| Age at death (years) | Cerebellum | 23 | 23 | 63.6 (17.1) | 70.9 (17.8) | 0.16 | Wilcoxon |
| Age at death (years) | Frontal | 23 | 22 | 63.6 (17.1) | 70.7 (18.2) | 0.184 | Wilcoxon |
| RIN | Cerebellum | 23 | 23 | 4.47 (1.51) | 5.70 (1.77) | 0.016* | Wilcoxon |
| RIN | Frontal | 23 | 22 | 6.29 (1.36) | 6.25 (0.971) | 0.917 | Wilcoxon |
| PMI (hours) | Cerebellum | 23 | 23 | 41.9 (24.3) | 55.2 (19.0) | 0.047* | Wilcoxon |
| PMI (hours) | Frontal | 23 | 22 | 41.9 (24.3) | 55.6 (19.4) | 0.044* | Wilcoxon |
| Sex (n Male: n Female) | Both | 46 | 45 | 12M: 11F | 13M: 10F | 1 | Chi-square |

### Supplementary Figures

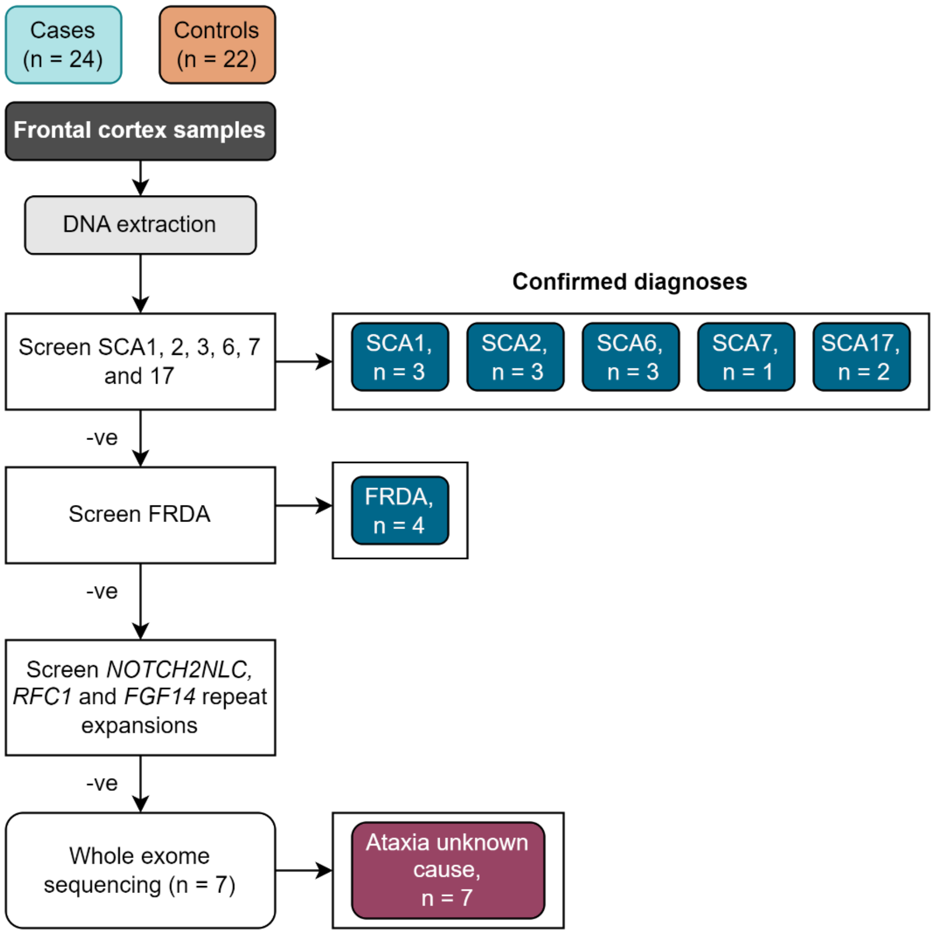


**Supplementary Figure 1. Workflow for molecular diagnosis of samples.** (**a**). DNA was extracted from all frontal cortex samples and screened for repeat expansions and allelic sizes for those loci estimated using repeat- prime polymerase chain reaction (RP-PCR). First, repeat expansions associated with spinocerebellar ataxia (SCA) 1, 2, 3, 6, 7 and 17 were screened for. Samples negative ("-ve") for these SCA expansions were subsequently screened for repeat expansions in *FXN* associated with Friedreich's ataxia (FRDA). All samples (including controls) that did not have a positive repeat expansion for any of the SCAs or FRDA screened were subsequently screened for repeats associated with recently described repeat expansion ataxia disorders. This included screening of *NOTCH2NLC* associated with neuronal intranuclear inclusion disease; *RFC1* associated with cerebellar ataxia, neuropathy, vestibular areflexia syndrome (CANVAS); and *FGF14* associated with SCA27B. Cases that were negative for these ten repeat expansion disorders were designated as having unknown molecular diagnosis. (**b**). Electropherograms illustrating examples of a positive repeat expansion in *ATXN1* with a typical sawtooth pattern and tailing (top panel) and electropherogran depicting no repeat expansion in the bottom panel.

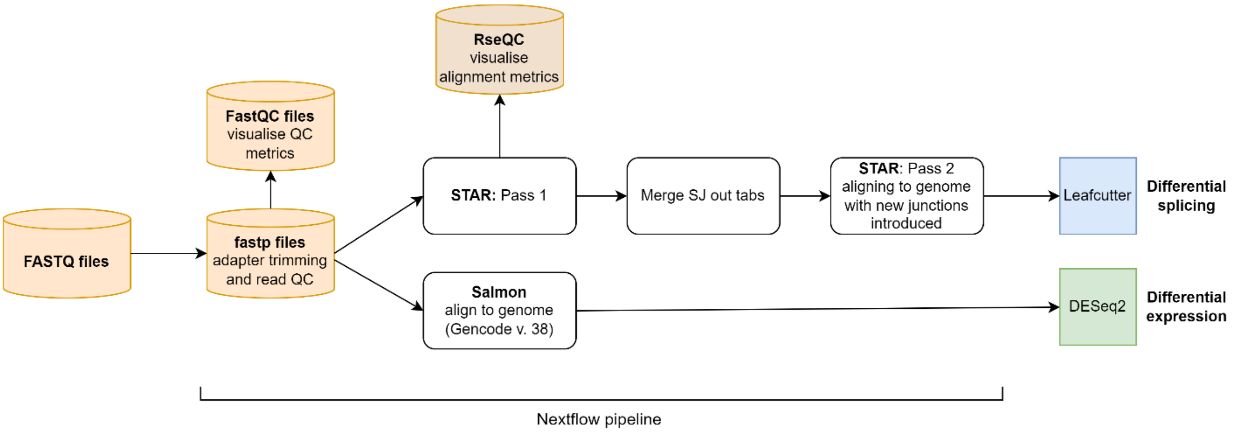


**Supplementary Figure 2. Workflow for data processing**. Workflow detailing pre-alignment quality control and alignment steps. Further data processing steps for input into Leafcutter and DESeq2 packages are also outlined. The quality control and alignment steps are included within a Nextflow pipeline.

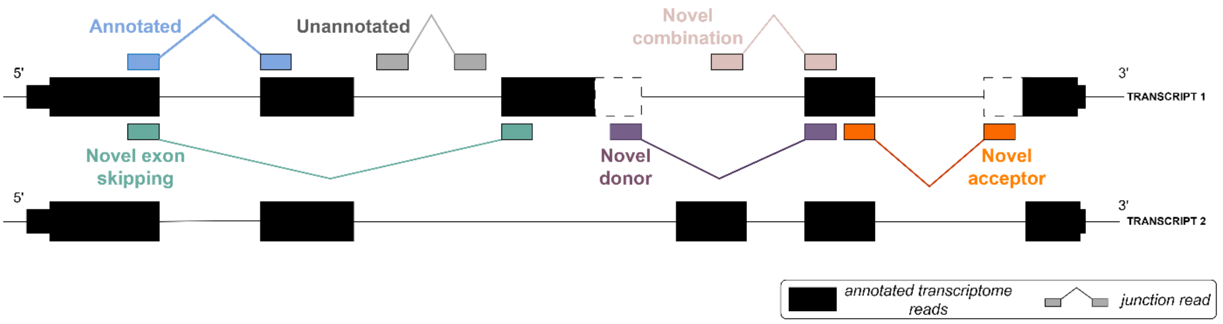


**Supplementary Figure 3. Splicing event annotation.** The annotated transcriptome (mapping to exons defined by Gencode v.38) is represented by the black filled boxes. Junction reads in relation to the existing annotation are also shown. Annotated junctions have both donor and acceptor splice sites that match the boundaries of an existing intron. Novel exon skip and novel combination junctions have donor and acceptor splice sites that overlap known exon boundaries but not of constitutive exons within the same transcript or different transcripts respectively. Novel donors (3' end) and novel acceptors (5' end) are junctions where only one end matches an existing exon boundary, and partially annotated to the reference transcriptome. Unannotated junctions have no overlaps with known exons. The two transcripts shown have shared and differing exons.

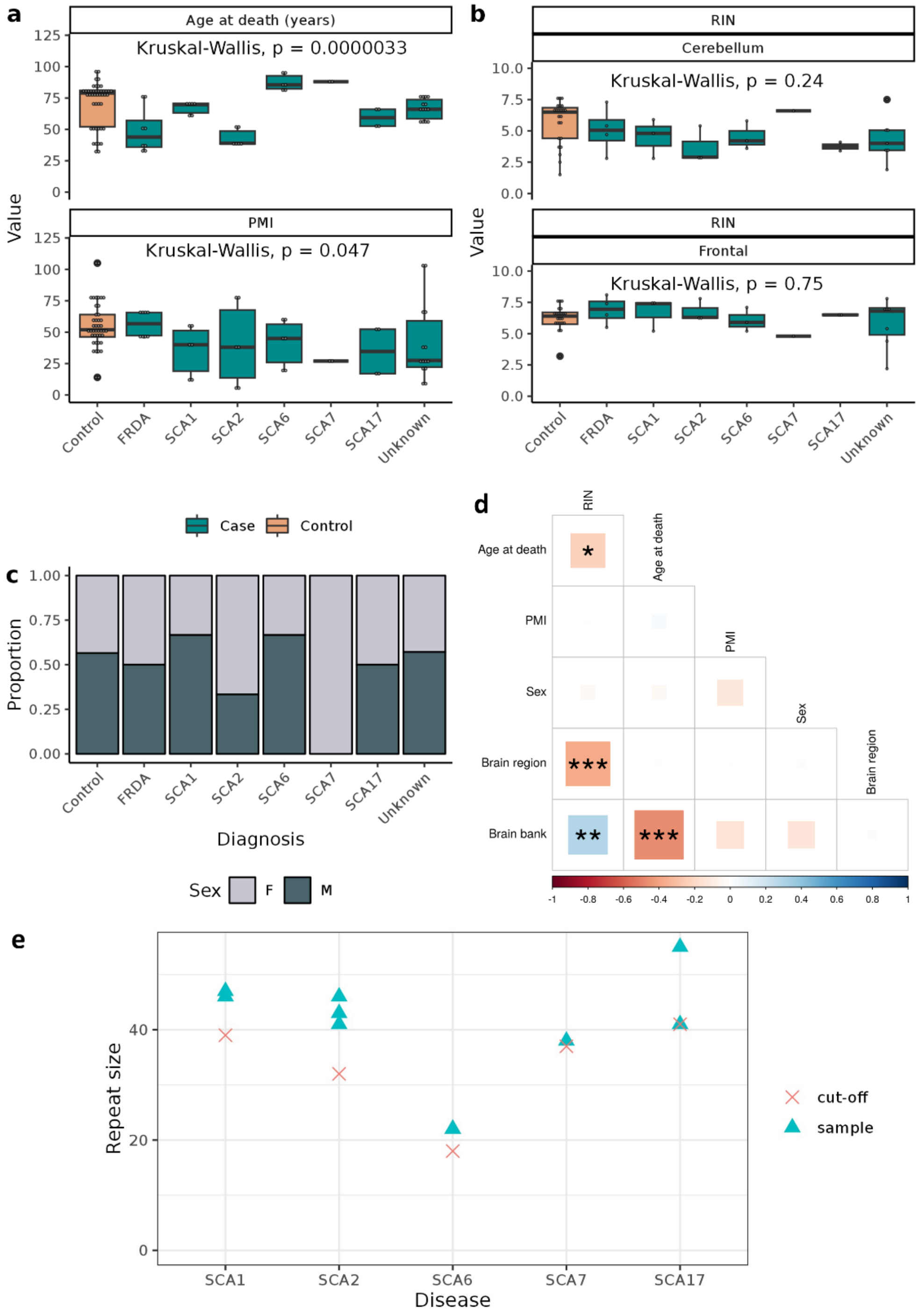


**Supplementary Figure 4. Comparison of sample measures and demographics.** Sample measures are compared across cases with and without (labelled as "unknown") molecular diagnoses. Age at death and postmortem interval (PMI) (**a**); RNA integrity number (RIN) across two brain regions (**b**) and proportion of female ("F") and male ("M") individuals (**c**) across the different diagnoses are shown. SCA represents spinocerebellar ataxia, FRDA represents Friedreich's ataxia and "unknown" refers to cases of ataxia without known molecular diagnosis. (**d**) Correlation plot showing the Pearson's correlation coefficient (scale from -1 to 1) between different sample metrics. Significance of the correlation are shown by superimposed asterixes: *, **, *** represent p-values of 0.01 – 0.05, 0.001 – 0.05 and < 0.001 respectively. Repeat expansion sizes of the SCAs were estimated using repeat prime PCR with the pathogenic repeat size threshold for repeat expansion for each of the diseases (defined by OMIM) are shown by the cross ("cut-off") (**e**).

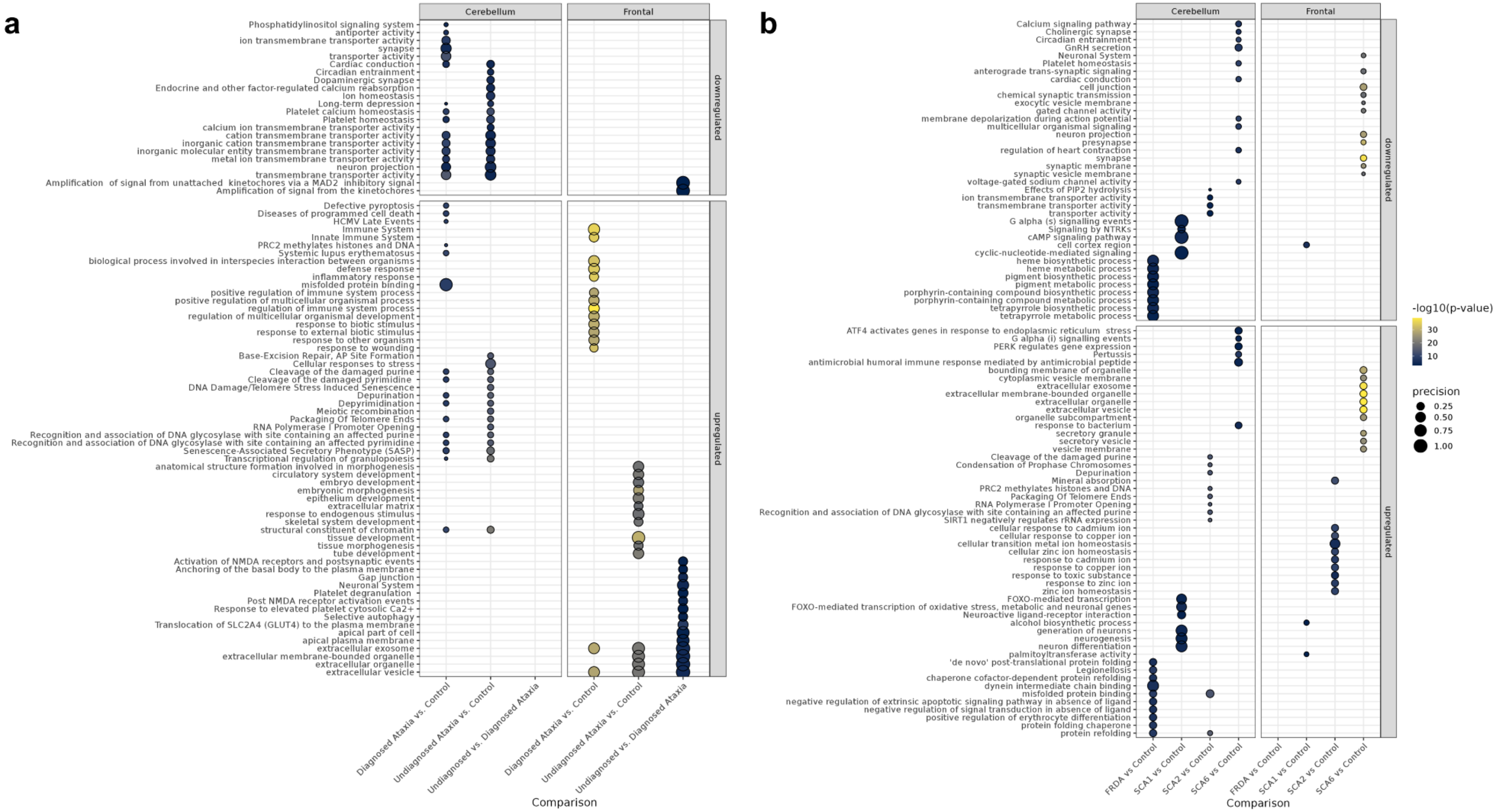

**Supplementary Figure 5. Pathway enrichment at Levels 2 and 3 of analysis.** Pathway enrichment analysis and significantly enriched gene ontology (GO) terms associated with Level 2 analysis, comparing molecularly diagnosed ("known") cases, molecularly undiagnosed ("unknown") cases and controls (**a**), and Level 3 analysis, comparing all cases with a molecular diagnosis in which n ≥ 3 (**b**).Only the most enriched pathways (defined by p value magnitude) are shown. Precision is from gprofiler package and defines the proportion of genes in the input gene list that can be assigned this term. P values given are gSCS-corrected.

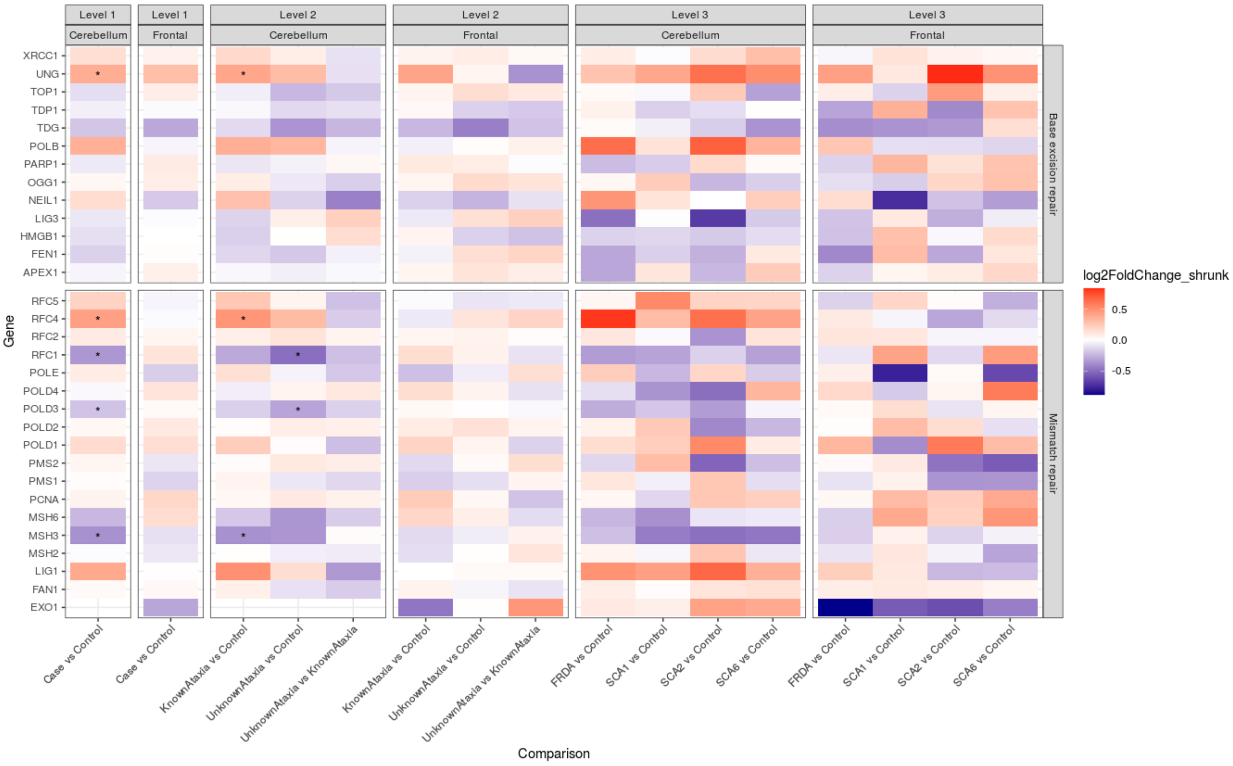


**Supplementary Figure 6. Differential expression of mismatch and base excision repair genes. * denotes significantly differentially expressed genes.**

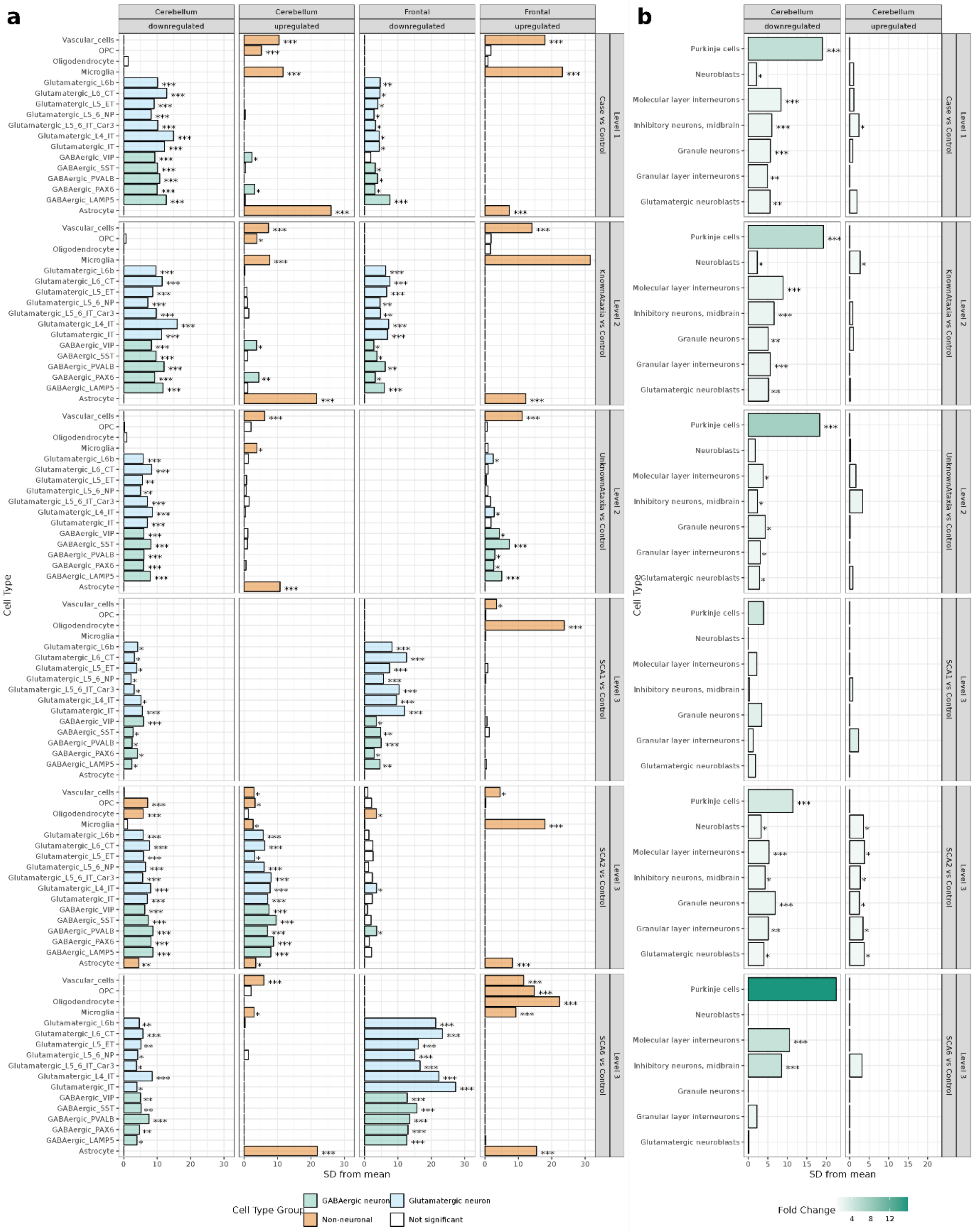


**Supplementary Figure 7. Expression-weighted cell type enrichment (EWCE) of differentially expressed genes.** EWCE of differentially expressed genes across the comparison groups within central nervous system cell types (**a**) and cerebellar-specific neuronal cell types (**b**). The standard deviations (SD) from the mean indicates the distance (in SD) of the target list of differentially expressed genes from the mean of the bootstrapped samples. The fold change denotes the change in expression of the target list compared to the bootstrapped list for a particular cell type. Significant results are indicated as follows: ***FDR $p < 0.001$; **FDR $p < 0.01$; *FDR $p < 0.05$. OPC stands for oligodendrocyte precursor cells. Description of cell types are presented in **Supplementary Table 3**.

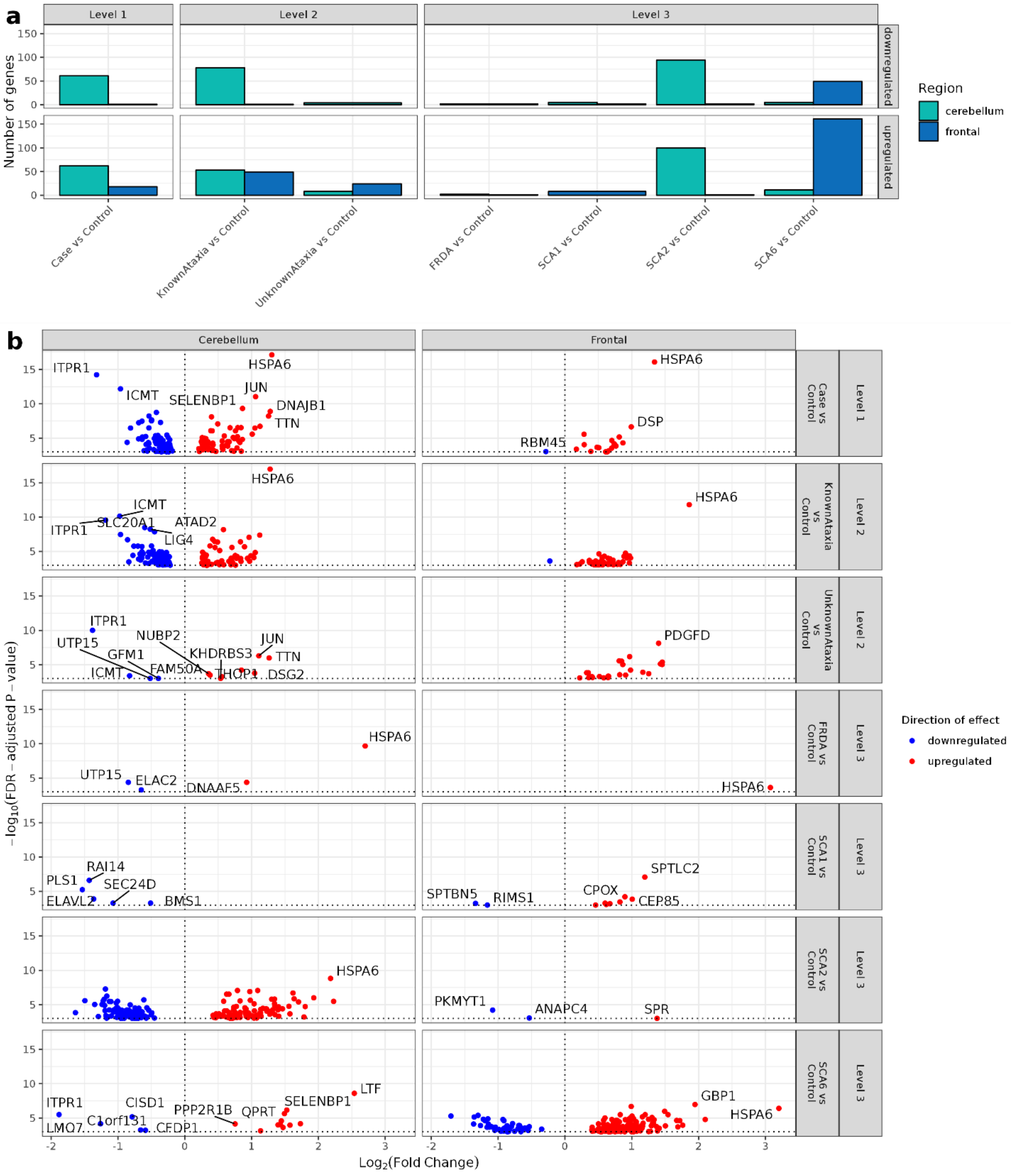


**Supplementary Figure 8. RNA-binding protein expression.** Number of genes associated with RNA-binding protein (RBP) function within differentially expressed genes at different levels of comparison (**a**). Differential expression of genes with RBP function (**b**).

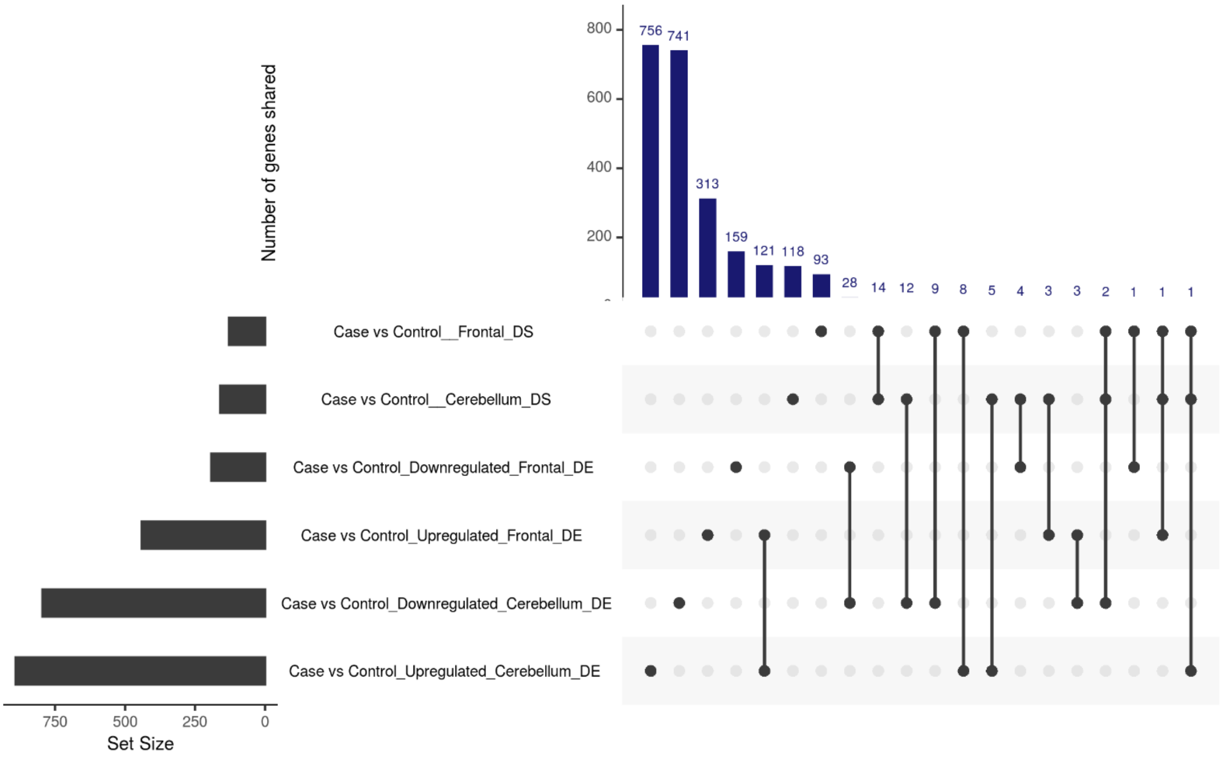


**Supplementary Figure 9. Number of overlapping differentially spliced and differentially expressed genes in Level 1 of analysis.** Comparison of the numbers of overlapping differentially spliced (DS) and differentially expressed (DE) genes across cases compared to controls

### Supplementary Methods

#### Protocol for diagnostic screening of common repeat expansion spinocerebellar ataxias

Tethering PCR was modified from the protocol described in Cagnoli *et al.* [reference]* by the NHS Neurogenetics Diagnostic Laboratory at Great Ormond Street Hospital. The PCR mastermix used for SCAs 1, 3 and 6 were the same: 2.5 mM dNTPs (Promega); 25 mM magnesium chloride; Flexi Buffer 5 X (Promega); Betaine 5M; GoTaq (5U/$\mu$l) (Promega); 100 ng/$\mu$l of DNA; 10 pmol/$\mu$l forward primer and 10 pmol/$\mu$l reverse primer. For SCA2 and SCA7, the same PCR mix was used with the exception of an additional DMSO (100%). Primers used are shown in Table C.2. For the repeat-prime PCR (RP-PCR), conditions were as follows (Table C.2). Following PCR, Liz 500 (ThermoFisher) mastermix made for capillary electrophoresis using 12$\mu$l HiDi formamide and 0.3$\mu$l of Liz 500 size standard per sample. This was mixed in the plate with PCR products for SCA 1, 2, 3, 6 and 7, 1 in 5 with nanopure water (ThermoFisher). The plate is then sealed and heated at 95 $^{\circ}$C for 3 minutes, then immediately cooled on ice for 3 minutes. This is then briefly spun down and loaded onto the ABI 3730 sequence analyser (ThermoFisher) using the fragment analysis protcol. CAG repeat number for each SCA corresponding to RP-PCR electropherogram peaks was calculated in GeneMapper using bins.

#### SCA17 screening

RP-PCR for SCA17 *TBP* expansion was carried out in a similar manner using fluorescent- labelled forward primers (Table C.1). PCR reagents and conditions were slightly altered compared to screening for SCAs 1, 2, 3, 6, 7 and 17. PCR mastermix comprised: 12.5 $\mu$l of AmpliTaq Gold 360 MasterMix (Thermofisher); 2.5 $\mu$l GC Enhancer; 5 pmol/$\mu$l of each primer; 1 $\mu$l DNA and Nanopure water up to 25 $\mu$l. PCR conditions were as follows: 95 $^{\circ}$C for 10 mins; 30 cycles of 95 $^{\circ}$C 30 seconds, 58 $^{\circ}$C for 30 seconds, 72 $^{\circ}$C for 30 seconds; 72 $^{\circ}$C for 7mins. The capillary electrophoresis and GeneMapper protocol was the same as for the other SCAs.

**Table C.1 PCR primers used.** Primers (F: forward; R: reverse) used for tethered repeat-prime PCR for SCAs 1, 2, 3, 6 and 7.

| Disease | Primer sequence |
|---|---|
| SCA1 | F: 5’-NED-TTTGCTGGAGGCCTATTCCACTCT-3’<br>R: 5’-GAGCCCTGCTGAGGTGCTGCTGCTGCTGCTG-3’ |
| SCA2 | F: 5’-VIC-TTTCGGCGGCTCCTTGGTCTC-3’<br>R: 5- AGCCGCGGGCGGCGGCTGCTGCTGCTGCTG-3’ |
| SCA3 | F: 5’-FAM-AGTCCAGTGACTACTTTGATTCG-3’<br>R: 5’- GTCCTGATAGGTCCCCCTGCTGCTGCTGCTG-3’ |
| SCA6 | F: 5’-VIC-TTTTTCCCCTGTGATCCGTAAGG-3’<br>R: 5’- CGGCCTGGCCACCGCCTGCTGCTGCTGCTG-3’ |
| SCA7 | F: 5’-FAM-TTTGAAAGAATGTCGGAGCGGG-3’<br>R: 5’-CTGCGGAGGCGGCGGCTGCTGCTGCTGCTG-3’ |
| SCA17 | F: 5’-FAM-GATGCCTTATGGCACTGGACTG-3’<br>R: 5’-CTGCTGGGACGTTGACTGCTG-3’ |

**Table C.2 PCR conditions.** Tethered repeat-prime PCR conditions for SCAs 1, 2, 3, 6 and 7.

| Temperature (°C) | Duration (minutes) | Cycles |
|---|---|---|
| 95 | 7 | x1 |
| 95<br>58 or 60 or 62<br>72 | 1<br>1<br>1 | x30-35 |
| 72<br>4<br>62 | 10<br>HOLD<br>SCA1 and 6 | x 30 |
| 60 | SCA 2 and 3 | X35 |
| 58 | SCA 7 | X35 |

#### FRDA screening

TP-PCR for FRDA *FXN* expansion was carried out in a similar manner. The three primers used were as follows:

- FATP-P1 5'-GCTGGGATTACAGGCGCGCGA-3';
- FATP-P3 FAM-5'-TACGCATCCCAGTTTGAGACG-3';
- FATP-P4 5'-TACGCATCCCAGTTTGAGACGGAAGAAGAAGAAGAAGAAGAA-3'.

PCR mastermix comprised: 10 $\mu$l of AmpliTaq Gold 360 MasterMix (Thermofisher); 2 $\mu$l GC Enhancer; 1 $\mu$l of 10 pmol/$\mu$l FATP-P3 (5'FAM) primer of each primer and $\mu$l primer mix: FATP-P1 10 pmol/$\mu$l + FATP-P4 1pmol/$\mu$l; 2 $\mu$l DNA and Nanopure water up to 25 $\mu$l. PCR conditions were as follows: 95 °C for 10 mins; 35 cycles of 95 °C for 1 min, 58 °C for 1 min, 72 °C for 1 min; 72 °C for 7 mins. The capillary electrophoresis and GeneMapper protocol was the same as for the SCAs.
